# Integrated multiomics analysis unveils how macrophages drive immune suppression in breast tumors and affect clinical outcomes

**DOI:** 10.1101/2024.11.09.622776

**Authors:** Youness Azimzade, Mads Haugland Haugen, Vessela Nedelcheva Kristensen, Arnoldo Frigessi, Alvaro Köhn-Luque

**Affiliations:** Oslo Center for Biostatistics and Epidemiology, University of Oslo, Oslo, Norway; Department of Tumor Biology, Institute for Cancer Research, Division of Cancer Medicine, Oslo University Hospital, The Norwegian Radium Hospital, Oslo, Norway; Department of Medical Genetics, Oslo University Hospital, University of Oslo, Oslo, Norway; Oslo Center for Biostatistics and Epidemiology, Oslo University Hospital, Oslo, Norway

## Abstract

Despite thorough characterizations of cellular compositions within the breast tumor microenvironment (TME), their implications for disease progression and patient prognosis are still poorly understood. Unraveling these effects is vital for identifying potential targets to improve treatment outcomes. In this study, we devised an explainable machine learning (XML) pipeline to scrutinize the associations between TME cellular constituents and relapse-free survival (RFS). By applying our pipeline to estimated cell fractions in the METABRIC and TCGA datasets and comparing these results with associations to pathological complete response (pCR) after neoadjuvant chemotherapy (NAC), we created a comprehensive catalog of the TME’s role based on 5000 patient samples. Our findings reveal an unexpected dichotomy in which macrophages correlate positively with pCR but negatively with RFS, particularly within estrogen receptor-positive (ER+) and Luminal A and B (LumA/B) cancer subtypes. We show that this pattern is driven by heterogeneity in breast tumors characterized by increasing levels of macrophage infiltration. Through imaging mass cytometry (IMC) analysis, we discovered that macrophages tend to accumulate in the vicinity of HLA-ABC^hi^ epithelial cells as their frequency increases in tumor tissues and also express elevated levels of HLA-ABC protein. Combining IMC with single-cell RNA sequencing (scRNA-seq) data, we uncovered a significant association between these HLA-ABC^hi^ macrophages and regulatory and exhausted T cells (T_Reg_ and T_Ex_), suggesting their involvement in immune suppression, likely by creating a chronically activated immunosuppressive TME. Subsequent cell-cell communication analysis predicted interactions between HLA-ABC^hi^ macrophages and T_Ex_ cells via the ligands SIGLEC9, ALCAM, and CSF1, and with T_Reg_ cells through APP, ANGPTL4, and SIGLEC9 signaling. Considering the clinical relevance of macrophages in ER+ (LumA/B) subtypes, our research enhances the characterization of macrophage-driven immune suppression in these tumors and identifies potential targets for immunomodulatory strategies.

## INTRODUCTION

The cellular composition of breast tumors has been of great interest for a long time [1], with the full extent of its diversity coming to light only in recent years due to advancements in scRNA-seq [2] and spatial omics data [3]. Despite having such a detailed overview, however, the contribution of cell types to tumor biology and clinical outcomes has remained less understood. This is partially because each of these data types captures certain aspects of cellular biology and bridging among them is inherently challenging. Clarifying the relative importance and involvement of cell types in clinical outcomes and the mechanisms by which they exert such effects can potentially identify new treatment targets and/or put current research on treatment into context.

Measuring gene expression of individual cells though scRNA-seq has provided a thorough characterization of the cellular composition of the breast TME [2, 4]. Additionally, by providing cell-type specific expression profiles, it has facilitated computational inspection of the abundance of cell types from previously analyzed samples–widely known as deconvolution [5]– providing the much needed statistical power for more comprehensive analysis [4, 6, 7].

Methodologies concentrating on spatial aspects of tumor samples have also significantly improved our understanding of the spatial configuration of breast cancer tissues among other cancer types [8]. The precise spatial context within tissue samples enables a comprehensive understanding of cellular and molecular interactions *in situ* [9]. Advancements in these methods, marked by enhanced imaging resolution, improved molecular detection sensitivity and sophisticated computational analysis methods, has enabled refined mapping of tissue samples, leading to a better understanding of spatial structures and interactions [3, 10, 11]. Integrating spatial omics with scRNA-seq data could further enhance our comprehension of cell-cell interactions, offering more detailed insights into the TME [12].

In the present paper, we explore the influence of cell types within TME on the RFS of breast cancer (BC) patients, by establishing a pipeline that employs XML to investigate this association. We introduce the “Survival Score”, a robust metric designed to quantitatively evaluate the association of various cell types with RFS. We calculate Survival Scores for cells across samples in the general population (GP) and tumor subtypes from the METABRIC (MBRC) and TCGA datasets. Juxtapose these freshly derived Survival Scores with the pCR Scores from our previous work [7], which evaluated the association of pCR after NAC, provides a comprehensive picture of the role of the TME in clinical outcome. Interestingly, it reveals that some cell types have an opposite association with RFS compared to pCR. Motivated by such observations for macrophages in the GP, ER+, LumA and LumB samples, we focus on them. We find that the persistent heterogeneity in breast tumors, characterized by varying macrophage infiltration levels, drives this behavior. To provide a functional understanding of the role of macrophages, we ran an integrated analysis of IMC and scRNA-seq data which revealed that a specific subset of macrophages, with distinct spatial distribution, are exclusively associated with immune suppressive TME. Our findings predict that this association can be a result of interactions between T_Ex_ and T_Reg_ with those macrophages through a few genes, identifying potential treatment targets in ER+ breast tumors.

## RESULTS

### Survival Scores

#### Cell types representing breast TME

To investigate the association between cell types and relapse-free survival, we first deconvolve bulk gene expression profiles (GEPs) and RNA-seq data using a signature matrix (SM). This is a cell-specific expression profile matrix, that is used as a reference to estimate cell fractions from bulk samples. Following the annotation in reference [4] with minor curations [7], the resulting SM includes 28 different cell types, including subsets of B cells, T cells, cancer-associated fibroblasts (CAFs), perivascular-like cells (PVLs), plasmablasts, myeloids, endothelial cells, normal epithelial cells and cancer cells.

#### Pipeline for Survival Scores

In each group of samples, existing data include clinical features such as age, tumor stage, tumor size and cell fractions calculated using deconvolution. To predict the risk of relapse, we used Random Survival Forests (RSF) [13] and Survival Support Vector Machines (SSVM) [14]. To assess the association of each cell type to survival probability, we first calculated their SHAP values [15] for all cell types. For the cell types that show a clear association between SHAP values and cell fractions (judged based on a linear fit; see Fig. 1), we checked if the association was similar across the two models. For such cell types, we calculated the average slope of the SHAP value versus cell fraction, normalized it and obtained the Survival Score. Finally, cell types were ranked based on their Survival Scores. As the final validation, we compared Survival Scores for MBRC and TCGA and excluded cell types that did not show similar scores.

**Figure 1.**
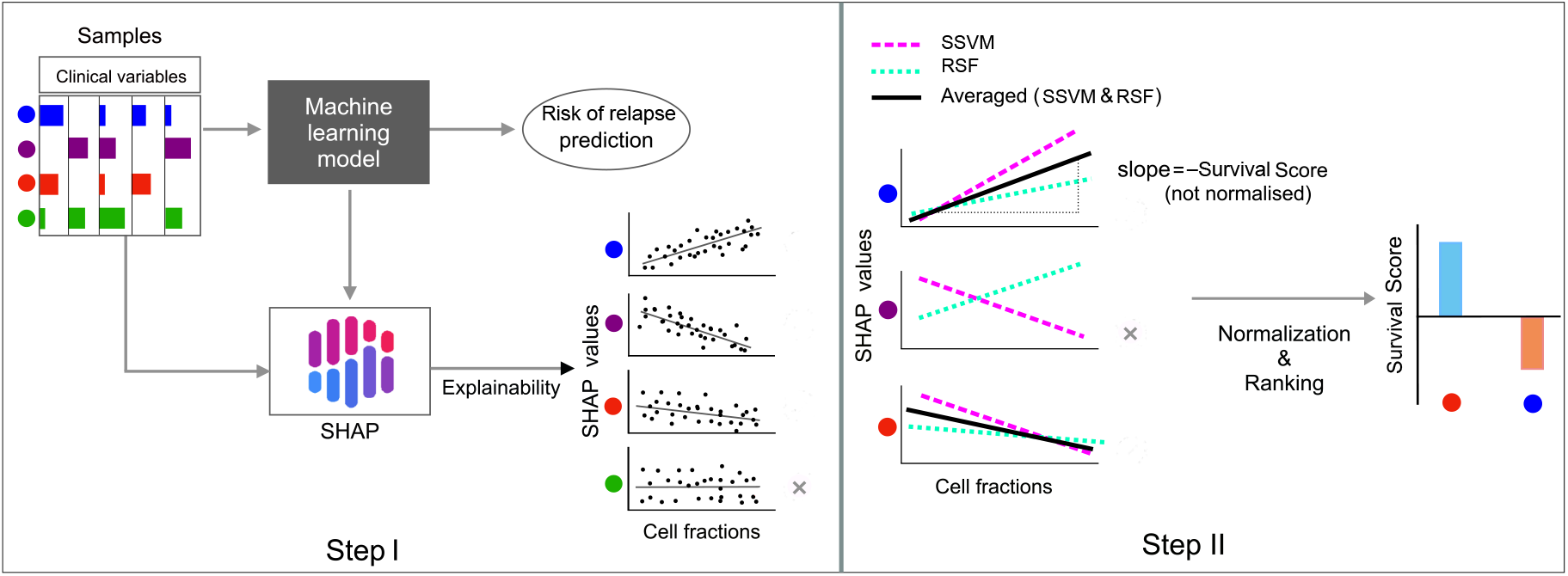
I) Cell fractions alongside available clinical variables (such as age, tumor stage and tumor size) are used to train an ML model that predicts the risk of relapse. SHAP values are calculated for all cell types in the given group of samples. Cell types that do not exhibit a clear association with SHAP values corresponding to the risk of relapse are excluded. II) For cell types with consistent associations in both models (either both negative or both positive), we calculate the average slope of the SHAP value versus cell fraction, normalize it and obtain the Survival Scores.

### Survival Scores in the GP

We apply our pipeline to the MBRC and TCGA datasets, separately. For each dataset, the pipeline identifies a wide range of cells associated with survival. We only discuss cell types that show a consistent association with survival across both datasets.

#### Cell types with positive Survival Scores

Among cancer cells, GenMod5 has positive Survival Scores in both datasets. Among immune cells, CD8 T cells, Natural Killer T (NKT) cells and Dendritic cells (DCs) are positively associated with survival. mature luminals also show a positive Survival Score.

#### Cell types with negative Survival Scores

GenMod6 and GenMod3 have negative Survival Scores. Among immune cells, natural killer (NK) cells and macrophages have negative Survival Scores. Among stromal cells, RGS5 endothelials, differentiated prevascular (PVL) cells and cancer-associated fibroblasts with myofibroblast characteristics (myCAFs) are noted. Interestingly, luminal progenitors also show a negative Survival Score.

Breast tumors include different subtypes with various cell types showing significantly different abundance among them (see Suppl. Fig. S1 and [7]). Additionally, each of these subtypes has undergone a different treatment plan [16] and has different survival rates [17]. Thus, the observed Survival Scores may reflect the combination of these differences. As such, we analyze these subtypes separately.

### Survival Scores in breast tumor subtypes, comparison to pCR Scores

We apply our pipeline to breast cancer subtypes, including ER subtypes (ER+/ER-) and PAM50 subtypes (LumA/LumB/Basal/HER2) [18]. For cell types that pass the model cross-validation at Step II and then show similar Survival Scores in both cohorts, we average over their scores in TCGA and MBRC to have a single Survival Score for each cell type (see Suppl. Figs S2 for Survival Scores in each cohort). For each cell type with a non-zero Survival Score in each group, we also include (if any) previously calculated pCR Scores [7] as shown in Fig. 3. As expected, most cell types that are associated with RFS are also associated with pCR, although the direction of the associations can be different.

**Figure 2.**
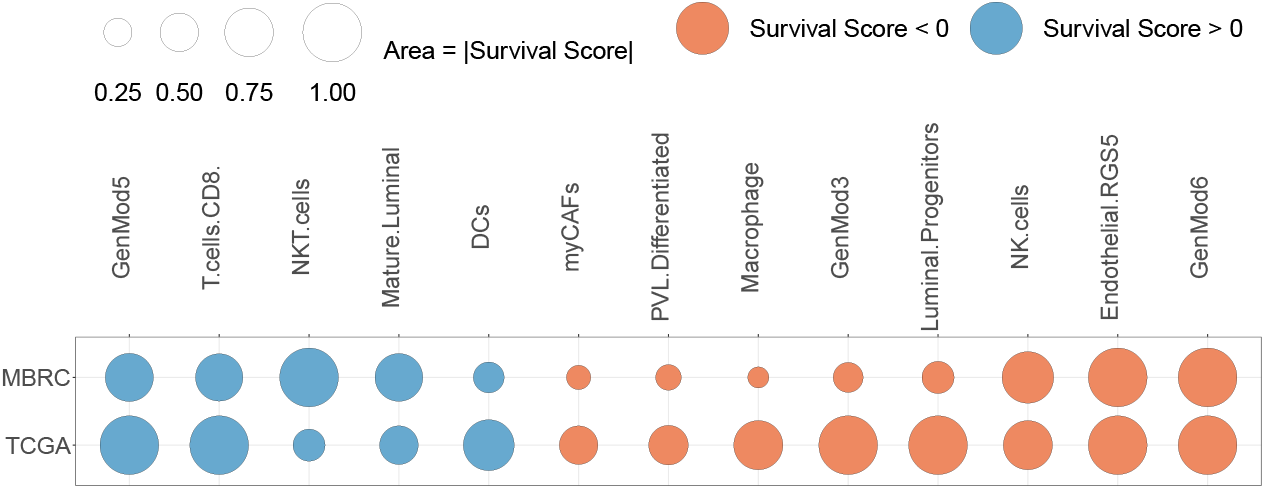
Ranked Survival Scores of cell types calculated in the GP of MBRC and TCGA datasets. A wide range of cell types, including cancer cells, immune cells and stromal cells consistently associated with survival across both datasets.

**Figure 3.**
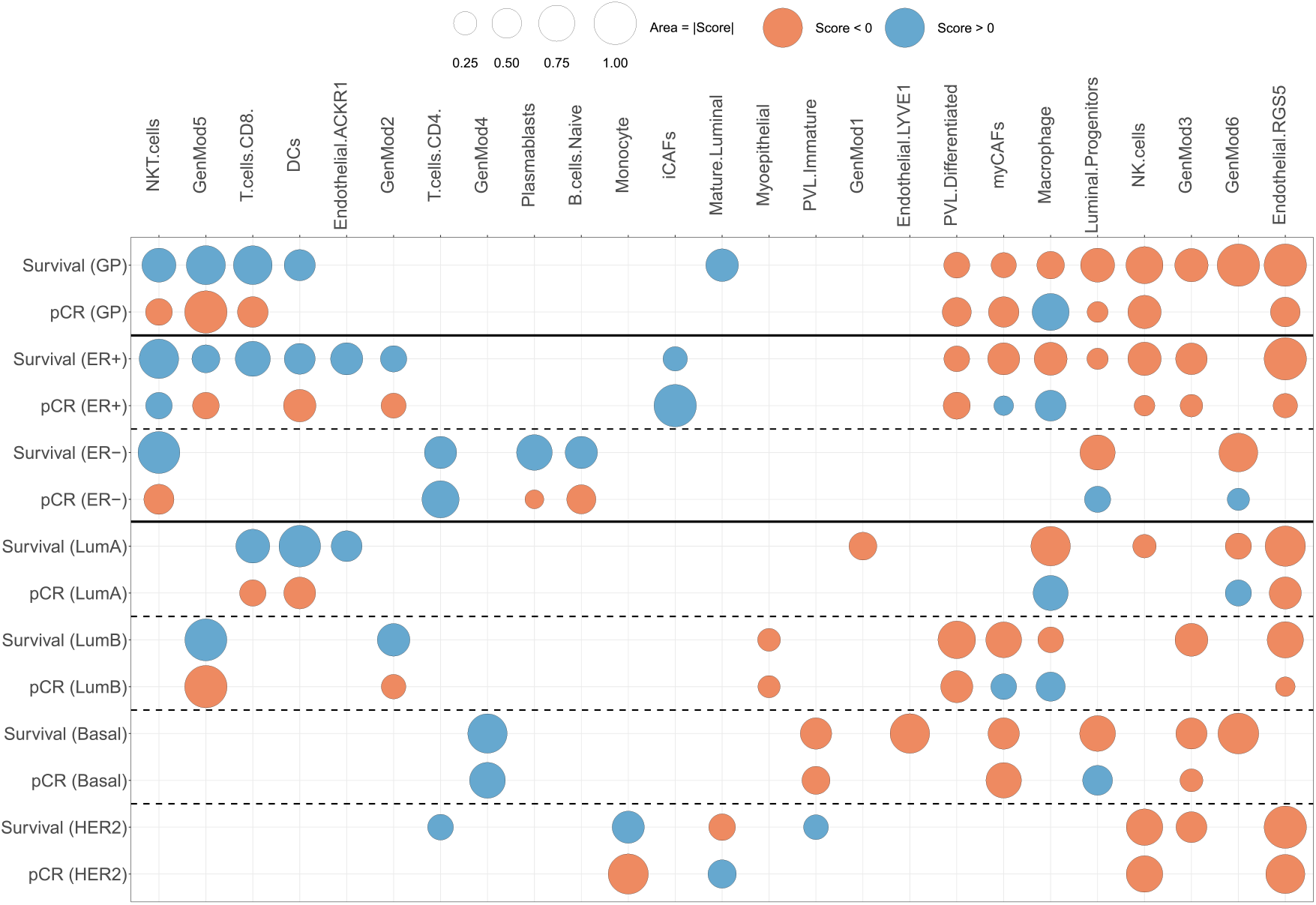
Comparison of pCR and Survival Scores in GP, ER and PAM50 Subtypes: While pCR and survival scores have the same sign for some cell types, for GenMod5 and macrophages, they consistently show opposite signs.

#### Cell types with similar pCR and Survival scores

Multiple cell types show aligned pCR and Survival scores, where both scores are either positive or negative, revealing that their association with the short-term and long-term outcomes of treatment are similar. RGS5 endothelials, GenMod3, NK cells, differentiated PVLs and myoepithelials with consistently negative scores and CD4 T cells, GenMod4 and iCAFs with consistently positive scores in at least some subtypes of tumors are examples of these cells.

#### Cell types with opposite scores

Within the GP, a variety of cell types show opposite scores. However, this trend shifts as we focus on ER and PAM50 subtypes, where we observe that most of these cell types disappear or their association no longer align with those found in the GP (see [7] for subtype-dependent roles of cell types). Two cell types, however, violate this pattern. GenMod5 exhibits a positive Survival score and negative pCR scores in three instances: GP, ER+ and LumB. Additionally, macrophages show negative Survival Scores and positive pCR Scores in four different instances: GP, ER+, LumA and LumB. In the following part, we explore different scenarios that could potentially explain such a scenario for macrophages (see SI and Suppl. Figs S3 for a quantitative assessment of alignment between pCR Scores and Survival Scores and analysis on GenMod5).

### Further exploration of Macrophages

#### Does macrophage molecular diversity drive the observed opposition in scores?

The molecular diversity in macrophages has the potential to drive this behavior as different subsets of macrophages can exhibit distinct associations with clinical outcome [19, 20]. Aside from the classical M1 and M2 categorization of macrophages, newer classifications based on scRNA-seq data have revealed relatively wider diversity. In our source scRNA dataset, which we used to create SMs, there are five molecular subsets named after their signature markers [4]: CXCL10 (M1-like), EGR1 (M2-like), SIGLEC1 (M2-like) and two Lipid-associated macrophages named LAM1-FABP5 and LAM2-APOE (see Fig. 4(a)).

**Figure 4.**
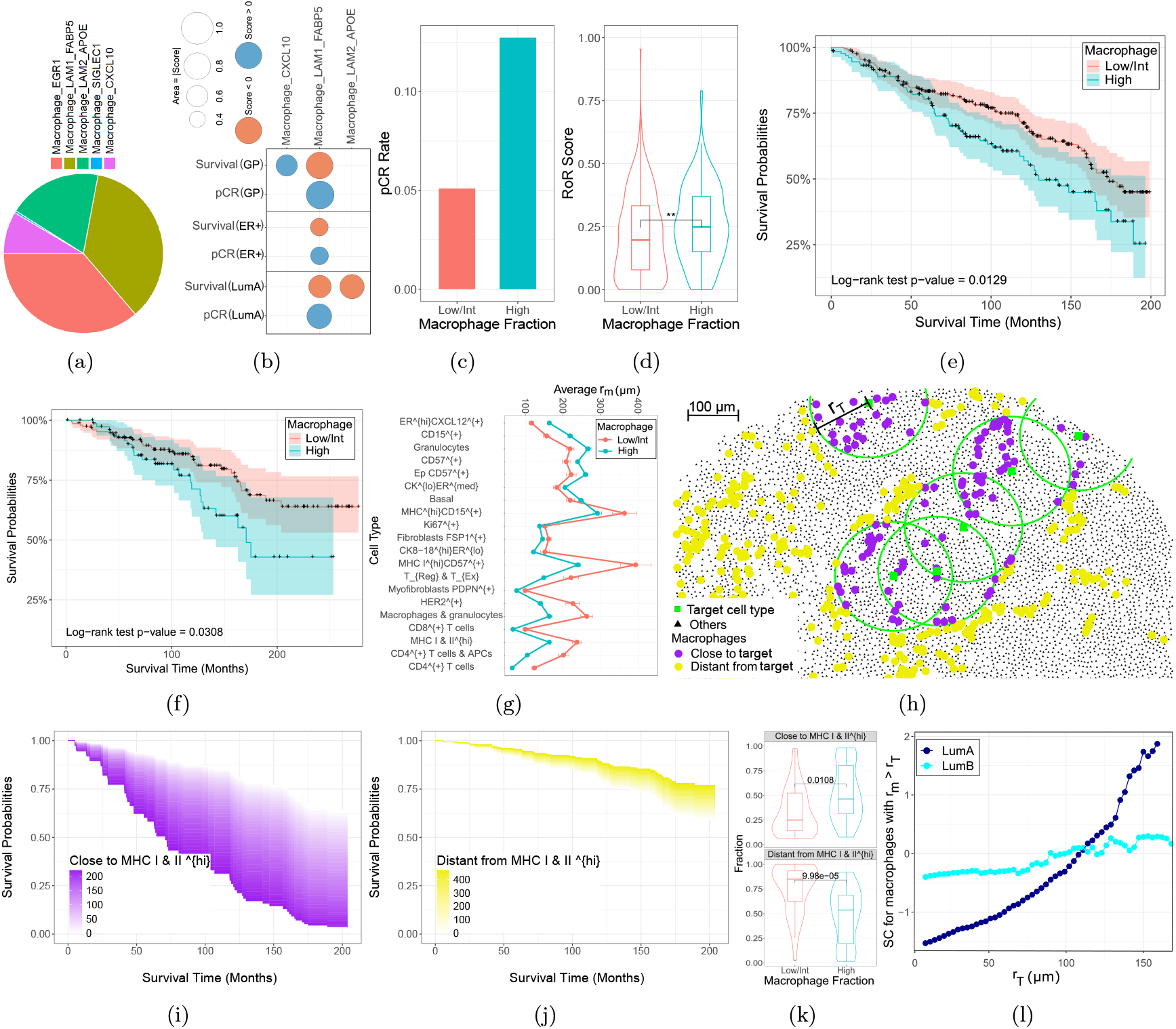
a) Subsets of macrophages with their corresponding proportions in scRNA-seq data. pCR and Survival Scores of subsets of macrophages in GP, ER+ and LumA. pCR rates (c) and RoR scores (d) compared between two groups of LumA (with high and Low/Int macrophage fractions) in the NAC cohort. KM plot for the two groups of LumA samples categorized based on the fraction of macrophages in deconvolution (e) IMC (f) data. g) Minimum distance, *r*_*m*_, between macrophages and other cell types from IMC data calculated and compared between the groups of lumA samples. h) In each sample, macrophages divided into two groups based on their *r*_*m*_ with respect to target cell type. Survival area plot for the effect of the frequency of macrophages close to MHC I & II ^hi^ (i) and distant from MHC I & II ^hi^ (j), revealing their opposite association with RFS. k) Fractions of two macrophage subsets in two groups of LumA samples. l) SC= −1 × (log hazard ratio) for macrophages with *r*_*m*_ *> r*_*T*_ in LumA and LumB samples.

We repeated the initial deconvolution step including the four most prevalent subsets of macrophages (CXCL10, EGR1, LAM1-FABP5 and LAM2-APOE) and then used it to estimate fractions of macrophage subsets (alongside other cell types mentioned before) in TCGA, MBRC and NAC datasets. Then, all data went through their corresponding pipelines and scores were calculated for all cell types. Fig. 4(b) shows the results for subsets of macrophages that passed the validation in GP, ER+, LumA and LumB (no subset passed the validation in LumB). CXCL12 (M1-like) macrophages exhibited positive Survival Scores in GP. LAM1, on the other hand, consistently showed negative Survival Scores in GP (in agreement with previous findings [4]), ER+ and LumA, while showing positive pCR scores in these groups. This observation reveals that while molecular diversity can lead to functional diversity of macrophages, even a subset of macrophages (here LAM1) can exhibit opposite pCR and Survival scores.

#### Macrophages infiltration stratifies LumA into subgroups with distinct clinical outcome

We investigated whether different subgroups of LumA could account for variations in pCR versus survival scores associated with macrophage levels. In our NAC cohort, we categorized LumA samples into two groups based on macrophage content: one with a low to intermediate fraction (67 % of samples) and another with a high fraction (33% of samples). These groups exhibited distinctly different clinical outcomes. The LumA group with a high macrophage fraction demonstrated a significantly higher pCR rate (see Fig. 4(c)). Concurrently, this group also faced a significantly elevated risk of recurrence (RoR) score [21] (refer to Fig. 4(d)). Furthermore, we analyzed LumA tumors from the MBRC dataset by dividing them similarly and assessing their survival rates. The group with a high macrophage content showed markedly lower survival probabilities, as illustrated in the Kaplan–Meier plot (see Fig. 4(e)). This pattern was consistent in ER+ and GP categories of both NAC and MBRIC datasets but was not observed in the TCGA dataset (see Suppl. Fig. S5).

#### Validation of heterogeneity in LumA tumors using spatial omics data

For a number of MBRC samples IMC data is available [22], allowing for the identification of macrophages. We select the LumA samples and calculate the fraction of macrophages in them by dividing their frequency by the number of all cells identified in each sample. We then divide the samples into two groups based on the fraction of macrophages and compare their survival probability. Interestingly, the group with a higher fraction of macrophages has a significantly lower survival probability, as shown in Fig. 4 (f), confirming our findings in the previous section using deconvolution data (the same behavior was observed for ER+ and GP, see Suppl. Fig. S6).

#### Distance from MHC I & II ^hi^ ECs reverses role of macrophages in RFS

After establishing that a higher fraction of macrophages is associated with lower survival probability, we intend to provide a functional understanding of their involvement in the TME. Considering the important role of the relative location of macrophages in the survival probability of esophageal cancer patients [23] and the general importance of distance in mathematical models of interactive populations [24, 25], we hypothesize that the vicinity of macrophages to other cell types may affect their functionality and, respectively, their role in clinical outcomes. Thus, we calculate the minimum distance, *r*_*m*_, between macrophages and other cell types in the IMC data. Interestingly, the comparison of *r*_*m*_ in two groups of LumA samples (with high or low/intense macrophages) shows that the majority of cell types (21 out of 31) exhibit significant differences, as shown in Fig. 4 (g), revealing that macrophages are in fact differently involved in the TME of these two groups.

Motivated by this observation, we divide macrophages into two groups based on their minimum distance to one target cell type (as shown in Fig. 4 (h)). In each sample, there will potentially be two subsets of macrophages, one close to the target cell type (with *r*_*m*_ *< r*_*T*_) and one distant from the target cell type (with *r*_*m*_ *> r*_*T*_); in samples with no target cell type or macrophages, both subsets are set to zero. The threshold distance, *r*_*T*_, is initially fixed as the median of the minimum distances, *r*_*m*_, between macrophages and the target cell type across LumA samples.

We then explored the association of each group of macrophages with RFS separately by fitting a Cox proportional hazards model [26]. Among all cell types, the vicinity of macrophages to epithelial cells with high MHC I & II levels (MHC I & II^hi^) seems to affect their role more drastically. As the survival area plot [27] in Fig. 4 (i) shows, a higher frequency of macrophages close to MHC I & II^hi^ leads to lower survival rates. On the other hand, a higher frequency of macrophages distant from MHC I & II^hi^ increases the survival probability, as shown in Fig. 4 (j). This observation was also validated in LumB and ER+ samples, where macrophages close to (and distant from) MHC I & II^hi^ show negative (and positive) effects on survival (see Suppl. Fig S7). It is worth mentioning that no other cell distance consistently reverses the role of macrophages, despite occasional observations of such effects. Additionally, as one might expect, the fraction of macrophages close to MHC I & II^hi^ is significantly higher in LumA samples with a high macrophage fraction, as shown in Fig. 4 (k).

#### Interaction length

While *r*_*T*_ was initially set as the median of *r*_*m*_ (equal to 132 *μm* for MHC I & II^hi^), it is not clear from what distance macrophages role in survival switches to positive. To explore this, we start with small values of *r*_*T*_ and explore the association of macrophages with *r*_*m*_ *> r*_*T*_ by fitting a Cox proportional hazards model. To be consistent with our previous section, we define the survival coefficient (SC) as −1 × (log hazard ratio). As Fig. 4 (l) shows, increasing *r*_*T*_ leads to a crossover in SC for both LumA and LumB samples at ∼ 110*μm*.

#### Macrophages close to MHC I & II^hi^ ECs are associated with an immune suppressive TME in LumA samples

To understand how proximity to MHC I & II^hi^ cells affects the functionality of macrophages, we compared the levels of markers between these two groups of macrophages. As Fig. 5 (a) shows, many markers, including antigen-presenting molecules HLA-ABC, show significant differences between the two groups of macrophages.

**Figure 5.**
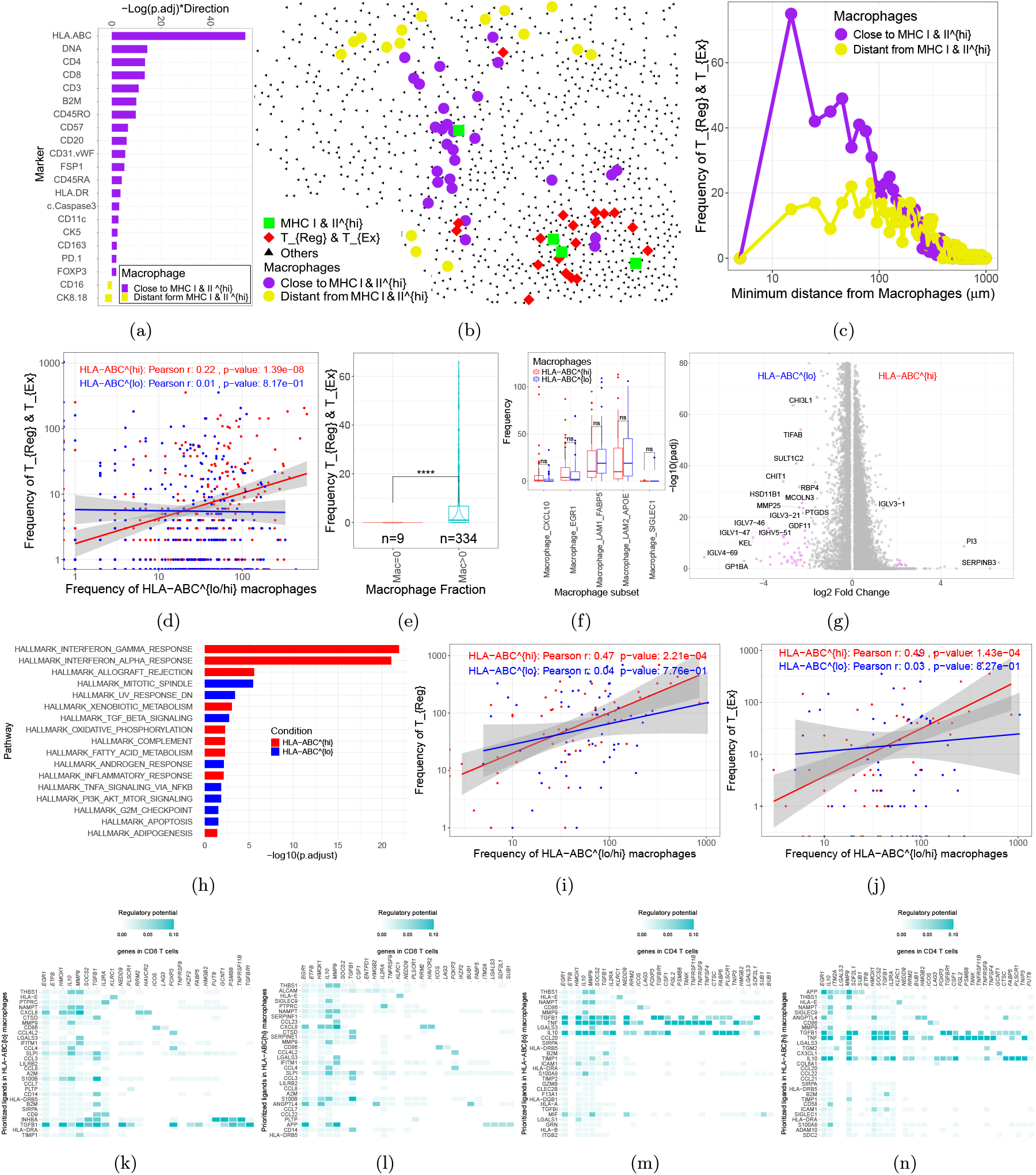
a) Comparison between levels of markers in two groups of macrophages in LumA samples. b) Spatial distribution of macrophages and T_Reg&_T_Ex_. c) Fraction of T_Reg&_T_Ex_ in vicinity of two groups of macrophages in LumA samples. d) Frequency of T_Reg_ vs frequency of HLA-ABC^lo^ and HLA-ABC^hi^ macrophages enumerated from spatial omics data for GP in breast tumors. e) Comparision between frequency of T_Reg&_T_Ex_ in samples with macrophages and without macrophages for samples including MHC I & II^hi^ cells. f) Comparison of frequency of HLA-ABC^lo^ and HLA-ABC^hi^ macrophages vs previously annotated subsets in scRNA seq data. DEA (g) and PEA (h) for HLA-ABC^lo^ vs. HLA-ABC^hi^ macrophages. i) Frequency of T_Reg_ vs two subsets of macrophages. j) Frequency of T_Ex_ vs two subsets of macrophages. Interactions between HLA-ABC^lo^ (k) and HLA-ABC^hi^ (l) macrophages and CD8 T cells. Interactions between HLA-ABC^lo^ (m) and HLA-ABC^hi^ (n) macrophages and CD4 T cells.

Macrophages with high levels of HLA-ABC are likely to be involved in immunosuppressive TME [28], which is typically characterized by the presence of exhausted T cells, T_Ex_ and regulatory T cells, T_Reg_, among other cell types [29]. The IMC dataset utilized thus far [22], identifies these two cell types, although they are collectively labeled as T_Reg&_T_Ex_. As Fig. 5 (b) shows, T_Reg&_T_Ex_ are positioned near macrophages close to MHC I & II^hi^ in one sample. Further analysis of T_Reg&_T_Ex_ frequency near two macrophage groups across LumA samples supports this finding (refer to Fig. 5(c)).

#### HLA-ABC^hi^ macrophages are exclusively associated with an immune suppressive TME in IMC data

Despite the insights the distance-based division of macrophages provided, this analysis is only applicable to samples with MHC I & II^hi^ ECs present. Hoping to define a more general and extendable definition of macrophages and inspired by Fig. 5 (a), we divide macrophages (43,000 cells from 700 samples) in all samples (including ER-samples) into two groups based on their HLA-ABC level: HLA-ABC^lo^ and HLA-ABC^hi^. We then explore how these groups are associated with T_Reg&_T_Ex_. As Fig. 5 (d) shows, the HLA-ABC^lo^ group is not associated with an immune suppressive TME. Interestingly, the HLA-ABC^hi^ group is significantly associated with T_Reg&_T_Ex_. It is worth mentioning that a partial correlation test accounting for the effect of MHC I & II^hi^ shows a significant correlation (p-value = 10^−8^). Additionally, in samples with MHC I & II^hi^ present, only those with macrophages also have T_Reg&_T_Ex_, as shown in Fig. 5 (e). A similar analysis on TNBC samples [11] for macrophages previously annotated as “M2 macrophages” confirms our finding for the exclusive association of HLA-ABC^hi^ macrophages with T_Ex_ (see Suppl. Fig. S8).

#### HLA-ABC^hi^ macrophages are exclusively associated with T_Reg_ and T_Ex_ in scRNA-seq data

Motivated by this observation in IMC data, we examined scRNA-seq data. We added a new dataset for scRNA-seq of breast cancer from [30]. We annotated T cells and myeloids separately by transferring the annotations from [4] using SEURAT [31]. Then, we divided all macrophages (10,700, across 57 samples) into two groups (HLA-ABC^lo^ and HLA-ABC^hi^). First, we explored whether HLA-ABC levels are associated with previous annotations of macrophages by comparing their abundance in these two groups. As Fig. 5 (f) shows, subsets of macrophages are evenly distributed between the two groups.

To compare these macrophages, we performed a differential expression analysis (DEA) between the two groups. Fig. 5 (g) shows the volcano plot for HLA-ABC^lo^ vs. HLA-ABC^hi^ macrophages, highlighting a variety of genes that are differentially expressed across these groups. We further conducted a pathway enrichment analysis (PEA) (see Fig. 5 (h)), which shows that HLA-ABC^hi^ macrophages are enriched in interferon response pathways, among others. Conversely, HLA-ABC^lo^ macrophages are enriched in the mitotic spindle, UV response DN and TGF beta signaling pathways.

Next, we explored the association between macrophages and T_Reg_ and T_Ex_ in these samples. The frequency of T_Reg_ does not show a significant association with HLA-ABC^lo^ macrophages but shows a significant positive association with HLA-ABC^hi^ cells, as shown in Fig. 5 (i). Similarly, the frequency of T_Ex_ shows a significant association only with HLA-ABC^hi^ macrophages, as Fig. 5 (j) shows. Previously, it was suggested that cancer cells with high levels of HLA-ABC are linked to immune-exhausted environments [12]. However, our results revealed that the association between HLA-ABC^hi^ macrophages and T_Reg_ and T_Ex_ is unique to HLA-ABC^hi^ macrophages (see Suppl. Fig. S9).

#### Interaction between HLA-ABC^hi^ macrophages and T_Ex_ cells

After establishing the exclusive association between HLA-ABC^hi^ vs. T_Ex_ and T_Reg_ cells, we aim to explore if this association stems from direct interactions. To explore this possibility, we utilize NicheNet [32], which is a cell-cell communication assay that predicts ligand-target links between interacting cells using scRNA-seq data. We then run NicheNet for both HLA-ABC^lo^ and HLA-ABC^hi^ macrophages and CD8 T cells. As Figs. 5 (k) and (l) show, various ligands from both groups of macrophages were predicted to regulate CD8 T cells. Interestingly, ligands with the highest regulation potential, including Activated Leukocyte Cell Adhesion Molecule (ALCAM) and Sialic Acid-binding Ig-like Lectin 9 (SIGLEC9) on macrophages, alongside genes such as Colony Stimulating Factor 1 (CSF1), were exclusively identified for HLA-ABC^hi^ macrophages/CD8 T cells. CSF1 is highly expressed by T_Ex_ cells in the TME and shapes myeloid cell recruitment and phenotype [33]. Further analysis suggests that CSF1, alongside other ligands on T_Ex_, regulates various genes on macrophages, including CCL20, CTSK and MMP9, which are involved in extracellular matrix remodeling (see Suppl. Fig. S10).

#### Interaction between HLA-ABC^hi^ macrophages and T_Reg_ cells

We performed a similar assay for the interaction between HLA-ABC^lo^ and HLA-ABC^hi^ macrophages and CD4 T cells. Interestingly, ligands such as Amyloid Precursor Protein (APP), Angiopoietin-like 4 (ANGPTL4) and SIGLEC9 exclusively appear in HLA-ABC^hi^ macrophages/CD4 T cells interactions (see Figs. 5 (m) and (n)).

## DISCUSSION

The analysis of the role of expression profiles in clinical outcomes has been of interest for a long time [34]. Exploiting new insight provided by high-resolution scRNA-seq data and XML, we provide a robust and reliable association between cellular components of TME and clinical outcomes based on more than 5000 samples. We showed that response to NAC and RFS for tumors are affected by different components of TME and even a certain cell type can have opposite association with these two outcomes. Such an observation stems from the fact that, in agreement with what has been established for ER+ and ER-samples [21, 35, 36], some groups of breast cancer patients can have good responses to NAC while having a higher risk of relapse. We showed that cell types such as GenMod5 and macrophages can discriminate samples into groups with different pCR rate/survival probabilities. Here we discuss cell types with high Survival Scores:

### NKT cells

As a subset of T cells sharing features of NK cells, NKT cells can exhibit both pro-and anti-tumor characteristics [37, 38]. Our results show a positive association between NKT cells and survival in the GP, as well as in ER+ and ER-subtypes.

### CD8 T cells

CD8 T cells have been extensively studied for their association with survival. In agreement with previous results [39], we find that higher fraction of CD8 T cells is associated with better survival in GP of BC as well as ER+ and LumA subtypes.

### DCs

DCs can contribute to immunosuppressive responses within the TME [40] or are positively associated with longer survival time [41] and can affect survival of breast tumors [42, 43]. Our results here show a positive role for DCs in survival in GP and in both ER+ and ER-subtypes.

### CD4 T cells

CD4 T cells, which can activate and regulate various aspects of innate and adaptive immunity and participate in tumor rejection [44]. CD4 T infiltration is shown to be positively associated with better survival in TNBC breast tumors [45, 46]. Our results also show a positive role for CD4 T cells in ER-and HER2 subtypes, confirming previous observations.

### differentiated PVLs

Resembling smooth muscle cells [47], differentiated PVLs demonstrate negative Survival Scores in the GP, ER+ and LumB. This is in general agreement with previous findings where the gene expression signature of vascular smooth muscle cells is associated with a poorer prognosis in cancer patients [48].

### myCAFs

myCAFs, enriched for myofibroblast pathways, can potentially affect the survival in breast tumors [49]. The direct association of myCAFs and survival, however, has not been explored. Our results show negative Survival scores in GP as well as in ER+, LumB and Basal subtypes.

### NK cells

Our results show negative Survival Scores for NK cells in GP as well as in ER+, LumA and HER2 subtypes. Combined with the observation of negative pCR Scores, their presence might be related to previously discovered intermediate immune infiltration with poor outcome [50].

### GenMod3

Enriched in EMT, IFN, Complement, GenMod3 shows negative Survival Scores in GP, ER+, LumB, Basal and HER2 subtypes. This finding may correspond with previous studies on the role of EMT enrichment in the loss of epithelial traits, the acquisition of mesenchymal characteristics and increased invasiveness [51].

### Endothelial RGS5

Our results show that Endothelial RGS5 shows high negative Survival and pCR scores across GP and various subtypes. RGS5, a pro-apoptotic/anti-proliferative protein, is a marker and regulatory factor in tumor angiogenesis, influencing vascular normalization, endothelial cell function and tumor growth. Expressed by the cancer vasculature triggered and retained by the proangiogenic microenvironment [52], it is associated with poor survival in various cancers [53]. While the frequency of endothelial cells is negatively associated with survival in colorectal cancer [54], observation of such a strong association should motivate further exploration of the role of Endothelial RGS5 in breast tumor tissues.

### GenMod5

As classical ER+ cells, GenMod5 can have higher resistance to chemotherapy, considering how estrogen levels affect the sensitivity of cells to chemotherapy [55]. On the other hand, ER+ tumors normally go through endocrine therapy [56] that aims to block estrogen’s ability to stimulate the growth of ER+ cancer cells. Therefore, it is somewhat expected that tumors with a higher fraction of GenMod5 respond better to endocrine therapy and demonstrate improved survival outcomes, despite having a poor response to NAC.

### Macrophages

Macrophages in tumor tissues, also referred to as tumor-associated macrophages (TAMs), play a critical role in tumor progression, angiogenesis and metastasis. Our result showed negative association between macrophages and survival in GP as well as ER+, LumA and LumB tumors. Interestingly, macrophages showed positive pCR scores in all of these groups. While molecular diversity in macrophages can lead to functional diversity [20] and different roles in clinical outcome [57], our findings revealed that even a specific subset of macrophages can display opposite associations with different clinical outcomes. As an alternative explanation, we showed that LumA/ER+/GP tumors can be divided into two subgroups with distinct pCR rate and survival probabilities based on the fraction of macrophages. Such a dominant role for macrophages is in line with recent observations on ER+ tumors where macrophages and not other immune cells were found to be the dominant immune cells in ER+ TME [58].

Exploring the dependency of the spatial location of macrophages and their role in RFS revealed that macrophages close to (distant from) ECs with high levels of MHC I & II show negative (positive) association with RFS. Furthermore, the composition of macrophages changes as the number of macrophages increases in tumor tissue. Tumors with higher infiltration of macrophages tend to have these cells located closer to MHC I & II^hi^ cells. Such macrophages are also colocalized with T_Reg_ and T_Ex_ in LumA samples and have higher levels of HLA-ABC. While spatial variation in macrophages has been observed in other cancer types [59], our findings reveal how the marker levels, spatial distribution and functionality of macrophages change as they accumulate in breast tumor tissues.

Our findings also reveal that macrophages can be divided based on their spatial distribution, with distinct associations with clinical outcomes and immunosuppression. This result is in line with gathering evidence on the role of the spatial location of macrophages on their functionality and phenotype [60–62].

Moreover, inspired by this classification, we used HLA-ABC level to divide macrophages into two groups which provided the opportunity to explore the role of macrophages in samples without MHC I & II cells as well as in scRNA-seq data. Interestingly, in IMC and scRNA-seq data, only HLA-ABC^hi^ macrophages (and not ECs) were associated with T_Reg_ and T_Ex_ and immunosuppressive TME, respectively. This observation challenges previous reports for breast tumors where HLA-ABC high ECs were reported to be associated with immunosuppression [12].

Considering the PEA results, HLA-ABC^hi^ macrophages are likely to contribute to immune exhaustion in the breast TME by creating a chronically activated, metabolically stressed and immunosuppressive environment [63]. HLA-ABC^hi^ macrophages are also enriched in OxPhos, suggesting that they are adapting to hypoxic TME, which enables them to engage in glycolysis, providing them with the required energy and biosynthetic resources to support their pro-tumoral functions [64].

PDL-1 high macrophages and T_Ex_ are previously shown to co-exist in high-grade ER- and ER+ breast tumors [65]. Considering the positive correlation between HLA-ABC and PDL1 levels (0.4 – see Suppl. Fig. S11 (a)), this prior observation might reflect the interaction we predict here. Additionally, macrophages with high levels of IRF8 also were suggested to drive T cell exhaustion [66]. Our PEA revealed that HLA-ABC^hi^ macrophages are enriched in IFN response pathways. Furthermore, analysis of HLA-ABC and IRF8 levels on macrophages shows a significant correlation (0.5) between the two markers (see Suppl. Fig. S11 (b)). These results collectively suggest that HLA-ABC level is associated with a variety of activities in macrophages and can be used as a robust measure to classify macrophages into groups with functional differences across different omics data.

Cell-cell communication assay identifies interaction axes unique to HLA-ABC^hi^ macrophages and T_Ex_ and T_Reg_, which can be potential treatment targets. In particular, SIGLEC9 ligand on HLA-ABC^hi^ macrophages was predicted to regulate genes related to immune suppression in both CD4 and CD8 cells. SIGLEC9 activity on macrophages can contribute to T cell exhaustion and, respectively, has gained interest as a target [67, 68]. Deletion of Siglece (murine homolog) resulted in dramatically restrained tumor development and prolonged survival in glioblastoma mouse models by activating both CD4+ T cells and CD8+ T cells through antigen presentation [69]. In breast cancer cell lines, targeting SIGLEC9 is shown to increase the cytotoxic activity of neutrophils [70]. Combined with our findings here, targeting SIGLEC9 presents a promising therapeutic strategy. By disrupting SIGLEC9 interactions, we can potentially enhance the immune response against tumors and improve the effectiveness of existing cancer treatments.

Additionally, ALCAM, also known as CD166, is exclusively the top ligand with the highest regulatory effect in HLA-ABC^hi^ macrophages regulating T_Ex_ cells. ALCAM is expressed on macrophages and can interact with its counter-receptor, CD6, found on T cells. This interaction is crucial for the physical and functional association between these cells. With ALCAM-based treatments under investigation [71, 72], our findings identify potential target patients benefiting from such treatments.

Cell-cell communication assay also identifies CSF1 as central to the interaction between T_Ex_ and HLA-ABC^hi^ macrophages, which are likely colocalized [33]. Reciprocally, CSF1 on T_Ex_ regulates a variety of genes on macrophages which are involved in inflammation and chemotaxis as well as extracellular matrix remodeling. This observation is in agreement with recent findings [33], where TAMs shown to interact with T_Ex_ in their vicinity. Our results confirm this observation and also reveals that only HLA-ABC^hi^ macrophages interact with T_Ex_ and contribute to immunosuppression. This finding is in line with recent results in ovarian cancer where co-localization of CD8 T cells with myeloids contributes to their exhaustion [73].

CSF1 has been of interest as a treatment target [74] and various CSF1-based drugs are under clinical trials [75] or concluded to be ineffective [76] in TNBC. Our results do not identify any association between macrophages in ER-or Basal-like tumors (two subsets more closely resembling TNBC) and clinical outcome, suggesting that the lack of response may be due to the minimal involvement of macrophages in these tumors rather than the ineffectiveness of the drug itself.

More generally, clinical trials on immunotherapies are concentrated on TNBC [76–78], ignoring their possible benefits in ER+ tumors, likely due to the old notion of ER+ tumors being immune “cold” and unresponsive. Our findings reveal the clinical relevance of macrophages in ER+ (LumA and LumB) subsets, which is in agreement with recent results [58]. This observation hints at the possible effect of immunotherapy in ER+ tumors. Furthermore, due to such a subtype-specific involvement of macrophages in clinical outcomes, the efficacy of macrophage-centered drugs in TNBC should not be extended to ER+ tumors and ER+ tumors should be explored in a separate trial of their own.

Altogether, our findings reveal the important role of macrophages in clinical outcomes, which is in line with a variety of results and current trends in macrophage-targeting immunotherapies [79–81]. Our work identifies subtypes of tumors where macrophages are involved in clinical outcomes and, respectively, macrophage-targeting immunotherapies are more likely to be effective. Finally, our results identify a few interaction axes as targets for macrophage-targeting immunotherapies. The clinical impact of macrophages is particularly significant in ER+ (LumA/B) breast tumors, where they are the dominant immune cells. This suggests that macrophage-targeting therapies might be more effective in these subtypes.

## METHODS

### Deconvolution

The scRNA-seq data source, including 100,000 cells, was obtained from 26 patients. Based on pseudobulk data analysis, this dataset comprises five LumA, three LumB, seven Basal-like, and four HER2-enriched samples, which can also be categorized as 11 ER+ and 15 ER-samples [4]. In line with our previous works [7, 82], this dataset was used as the reference for cell type-specific expression profiles.

#### Bulk Datasets

We used TCGA for breast cancer and the MBRC dataset as our two main datasets for RFS analysis. Bulk RNA-seq data of TCGA and GEPs data of MBRC were downloaded from cBioPortal. TCGA data were log2-normalized and scaled, while METABRIC data were scaled. We estimated cell fractions using either the “Impute Cell Fractions” module of the CIBERSORTx webtool with default settings or the Docker version of CIBERSORTx with similar settings. Our previously created 10 SMs [7] were used as references. We then averaged over all 10 estimates to increase the reproducibility of our results. Additionally, the following datasets were used in the NAC cohort: E-MTAB-4439 [83], GSE18728 [84], GSE19697 [85], GSE20194 [86], GSE20271 [87], GSE22093 [88, 89], GSE22358 [90], GSE42822 [91], GSE22513 [92], GSE25066 [93], GSE32603 [94], GSE32646 [95], GSE37946 [96], GSE50948 [97], GSE23988 [88].

### Machine Learning

Through the application of deconvolution, we were able to estimate the proportions of 28 distinct cell types, while also computing the ratios of prominent cell categories, namely B cells, T cells, CAFs, PVLs, myeloids, endothelial cells, normal epithelial cells, and cancer cells. Accompanying these cellular fractions are two supplementary attributes: ER status and PAM50 subtypes [21]. In our preprocessing stage, we designated positive (negative) ER statuses as 1 (0), respectively. Furthermore, we introduced dummy variables to represent the various PAM50 subtypes, transforming the single PAM50 subtype feature into five distinct features.

#### ML Models

We developed two prediction models: Random Survival Forests (RSF) and Survival Support Vector Machines (SSVM), utilizing the Scikit-Learn library [98] within the Python programming language. For model evaluation, we used the C-index. Hyperparameter tuning was performed using GridSearchCV. The best SSVM and RSF showed C-indexes of 0.683 (±0.012) and 0.686 (± 0.011) in METABRIC and 0.731 (±0.021) and 0.676 (± 0.041) in TCGA using 5-fold cross-validation, respectively.

#### SHAP Values

We utilized SHAP (SHapley Additive exPlanations) [15], a post-hoc explanation method that calculates the marginal feature contribution to the prediction per sample and then estimates the role of each cell population in the risk of relapse across the population.

To establish a measure for the influence of a cell type fraction, we conducted a linear regression analysis between SHAP values and cell type fractions. By evaluating the slope of this regression line along with its corresponding confidence interval, we assessed the overall association of the cell type fraction with the risk of relapse in each model (SSVM and RSF).

#### Survival Scores

For cell types that showed a similar association with the risk of relapse in both models (SSVM and RSF), we proceeded to calculate the Survival Score in their corresponding cohort. This included an initial normalization of all cell type fractions to rectify their varying abundances. Additionally, SHAP values were scaled across all cell types, providing a comparable scale for their values in both ML models. Next, a linear regression model was fitted to the SHAP values from both models plotted against their respective fractions.

Following this, the slopes of the fitted regression line were normalized and multiplied by − 1, assigning them the label of Survival Score. A positive (negative) Survival Score indicates a corresponding positive (negative) correlation between the fraction of a particular cell type and the likelihood of having a longer RFS.

#### Validation

We first performed in-cohort cross-validation of models that compared SHAP values predicted by two models, and only cell types with similar associations based on both models passed this stage. The second stage of validation was carried out across cohorts, and only cell types with similar Survival Scores passed this stage. Then, cell types were ordered based on their Survival Score across both cohorts, as illustrated in Fig. 2. For these cell types, we also averaged the Survival Scores to compare with pCR Scores, as shown in Fig. 3.

### Spatial Omics

IMC data from [22] were downloaded from zenodo.org/5850952, and IMC data for the TNBC cohort from [11] were downloaded from zenodo.org/7990870. Cells were already phenotyped. Distances between cells were calculated using the proxy function [99] in R.

### Survival Analysis

The association of cell frequencies and RFS was explored using the Survival package [100] in R. KM plots were generated using the survfit function. To perform a log-rank test to compare survival curves, the survdiff function was employed.

### scRNA-seq Data

In this study, two scRNA-seq datasets were used. scRNA-seq data from [4] were downloaded from broadinstitute.org/SCP1039. Counts and coarse-grained annotations from [30] were downloaded from lambrechtslab.sites.vib.be. The annotations for each major cell type from [4] were then transferred to [30] using Seurat [31]. For each major cell type, the common genes in both datasets were first identified. Next, anchors were defined using the FindTransferAnchors function, and cell phenotypes were predicted using the TransferData function.

### Cell-Cell Communication Assay

To run NicheNet [32], we followed the ligand activity geneset vignette. To run such an analysis, scRNA-seq data of candidate cell types and genes likely involved in the interaction are required. In the interaction between macrophages and T cells, HLA-ABC^lo^ (^hi^) macrophages were set as the sending cells, while CD4 and CD8 T cells were set as the receiving cells. For the sending cells, all genes expressed in these cells were selected as genes of interest. For the receiving cells, we selected 42 genes associated with T cell exhaustion [101] (such as HAVCR2, FABP5, CSF1, PDCD1, and LAG3), along with 31 other genes involved in regulatory T cells (such as FOXP3, IL2RA, CTLA4, GITR, and EGR1).

## Code availability

Codes are available at https://github.com/YounessAzimzade/Survival-XML-TME-BC.

## ACKNOWLEDGEMENTS

We would like to thank Parastoo Shahrouzi and Mads Haugen for reading the manuscript and their feedback. This project received funding from the European Union’s Horizon 2020 Research and Innovation Program under Grant Agreement No. 847912, BigInsight (NFR project 237718) and Integreat - The Norwegian center for knowledge-driven machine learning (NFR project 332645).

## References

[1] M. Allinen, R. Beroukhim, L. Cai, C. Brennan, J. Lahti-Domenici, H. Huang, D. Porter, M. Hu, L. Chin, A. Richardson, et al., Molecular characterization of the tumor microenvironment in breast cancer, Cancer cell 6, 17 (2004).

[2] E. Azizi, A. J. Carr, G. Plitas, A. E. Cornish, C. Konopacki, S. Prabhakaran, J. Nainys, K. Wu, V. Kiseliovas, M. Setty, et al., Single-cell map of diverse immune phenotypes in the breast tumor microenvironment, Cell 174, 1293 (2018).

[3] H. W. Jackson, J. R. Fischer, V. R. Zanotelli, H. R. Ali, R. Mechera, S. D. Soysal, H. Moch, S. Muenst, Z. Varga, W. P. Weber, et al., The single-cell pathology landscape of breast cancer, Nature 578, 615 (2020).

[4] S. Z. Wu, G. Al-Eryani, D. L. Roden, S. Junankar, K. Harvey, A. Andersson, A. Thennavan, C. Wang, J. R. Torpy, N. Bartonicek, et al., A single-cell and spatially resolved atlas of human breast cancers, Nature Genetics 53, 1334 (2021).

[5] Y. Im and Y. Kim, A comprehensive overview of rna deconvolution methods and their application, Molecules and Cells 46, 99 (2023).

[6] L. Kester, D. Seinstra, A. G. van Rossum, C. Vennin, M. Hoogstraat, D. van der Velden, M. Opdam, E. van Werkhoven, K. Hahn, I. Nederlof, et al., Differential survival and therapy benefit of patients with breast cancer are characterized by distinct epithelial and immune cell microenvironments, Clinical Cancer Research 28, 960 (2022).

[7] Y. Azimzade, M. Haugen, X. Tekpli, C. B. Steen, T. Fleischer, D. Kilburn, H. Ma, E. V. Egeland, G. Mills, O. Engebraaten, et al., Explainable machine learning reveals the role of the breast tumor microenvironment in neoadjuvant chemotherapy outcome, bioRxiv, 2023 (2023).

[8] V. Marx, Method of the year: spatially resolved transcriptomics, Nature Methods 18, 9 (2021).

[9] H. R. Ali and R. B. West, Spatial biology of breast cancer, Cold Spring Harbor Perspectives in Medicine, a041335 (2023).

[10] H. R. Ali, H. W. Jackson, V. R. Zanotelli, E. Danenberg, J. R. Fischer, H. Bardwell, E. Provenzano, O. M. Rueda, S.-F. Chin, et al., Imaging mass cytometry and multiplatform genomics define the phenogenomic landscape of breast cancer, Nature Cancer 1, 163 (2020).

[11] X. Q. Wang, E. Danenberg, C.-S. Huang, D. Egle, M. Callari, B. Bermejo, M. Dugo, C. Zamagni, M. Thill, A. Anton, et al., Spatial predictors of immunotherapy response in triple-negative breast cancer, Nature 621, 868 (2023).

[12] S. Tietscher, J. Wagner, T. Anzeneder, C. Langwieder, M. Rees, B. Sobottka, N. de Souza, and B. Bodenmiller, A comprehensive single-cell map of t cell exhaustion-associated immune environments in human breast cancer, Nature Communications 14, 98 (2023).

[13] H. Ishwaran, U. B. Kogalur, E. H. Blackstone, and M. S. Lauer, Random survival forests, (2008).

[14] V. Van Belle, K. Pelckmans, J. A. Suykens, and S. Van Huffel, Survival svm: a practical scalable algorithm., in ESANN (2008) pp. 89–94.

[15] S. M. Lundberg and S.-I. Lee, A unified approach to interpreting model predictions, Advances in Neural Information Processing Systems 30 (2017).

[16] A. G. Waks and E. P. Winer, Breast cancer treatment: a review, Jama 321, 288 (2019).

[17] E. Orrantia-Borunda, P. Anchondo-Nuñez, L. E. Acuña-Aguilar, F. O. Gómez-Valles, and C. A. Ramírez-Valdespino, Subtypes of breast cancer, Breast Cancer [Internet] (2022).

[18] C. M. Perou, T. Sørlie, M. B. Eisen, M. Van De Rijn, S. S. Jeffrey, C. A. Rees, J. R. Pollack, D. T. Ross, H. Johnsen, L. A. Akslen, et al., Molecular portraits of human breast tumours, Nature 406, 747 (2000).

[19] C. D. Mills, K. Kincaid, J. M. Alt, M. J. Heilman, and A. M. Hill, M-1/m-2 macrophages and the th1/th2 paradigm, The Journal of immunology 164, 6166 (2000).

[20] P. J. Murray and T. A. Wynn, Protective and pathogenic functions of macrophage subsets, Nature Reviews Immunology 11, 723 (2011).

[21] J. S. Parker, M. Mullins, M. C. Cheang, S. Leung, D. Voduc, T. Vickery, S. Davies, C. Fauron, X. He, Z. Hu, et al., Supervised risk predictor of breast cancer based on intrinsic subtypes, Journal of Clinical Oncology 27, 1160 (2009).

[22] E. Danenberg, H. Bardwell, V. R. Zanotelli, E. Provenzano, S.-F. Chin, O. M. Rueda, A. Green, E. Rakha, S. Aparicio, O. Ellis, et al., Breast tumor microenvironment structures are associated with genomic features and clinical outcome, Nature Genetics 54, 660 (2022).

[23] X. Ma, Z. Guo, X. Wei, G. Zhao, D. Han, T. Zhang, X. Chen, F. Cao, J. Dong, L. Zhao, et al., Spatial distribution and predictive significance of dendritic cells and macrophages in esophageal cancer treated with combined chemoradiotherapy and pd-1 blockade, Frontiers in Immunology 12, 786429 (2022).

[24] Y. Azimzade and A. Mashaghi, Search efficiency of biased migration towards stationary or moving targets in heterogeneously structured environments, Physical Review E 96, 062415 (2017).

[25] Y. Azimzade, Dynamics of motile populations in heterogeneous environments, arXiv preprint arXiv:1710.07937 10.48550/arXiv.1710.07937 (2017).

[26] J. Fox and S. Weisberg, Cox proportional-hazards regression for survival data, An R and S-PLUS companion to applied regression 2002 (2002).

[27] R. Denz and N. Timmesfeld, Visualizing the (causal) effect of a continuous variable on a time-to-event outcome, Epidemiology 34, 652 (2023).

[28] R.-Y. Ma, A. Black, and B.-Z. Qian, Macrophage diversity in cancer revisited in the era of single-cell omics, Trends in Immunology (2022).

[29] Y. Tie, F. Tang, Y.-q. Wei, and X.-w. Wei, Immunosuppressive cells in cancer: mechanisms and potential therapeutic targets, Journal of Hematology & Oncology 15, 61 (2022).

[30] A. Bassez, H. Vos, L. Van Dyck, G. Floris, I. Arijs, C. Desmedt, B. Boeckx, M. Vanden Bempt, I. Nevelsteen, K. Lambein, et al., A single-cell map of intratumoral changes during anti-pd1 treatment of patients with breast cancer, Nature Medicine 27, 820 (2021).

[31] Y. Hao, T. Stuart, M. H. Kowalski, S. Choudhary, P. Hoffman, A. Hartman, A. Srivastava, G. Molla, S. Madad, C. Fernandez-Granda, and R. Satija, Dictionary learning for integrative, multimodal and scalable single-cell analysis, Nature Biotechnology 10.1038/s41587-023-01767-y (2023).

[32] R. Browaeys, W. Saelens, and Y. Saeys, Nichenet: modeling intercellular communication by linking ligands to target genes, Nature Methods 17, 159 (2020).

[33] K. Kersten, K. H. Hu, A. J. Combes, B. Samad, T. Harwin, A. Ray, A. A. Rao, E. Cai, K. Marchuk, J. Artichoker, et al., Spatiotemporal co-dependency between macrophages and exhausted cd8+ t cells in cancer, Cancer Cell 40, 624 (2022).

[34] H. M. Bøvelstad, S. Nygård, H. L. Størvold, M. Aldrin, Ø. Borgan, A. Frigessi, and O. C. Lingjærde, Predicting survival from microarray data—a comparative study, Bioinformatics 23, 2080 (2007).

[35] H. G. Russnes, O. C. Lingjærde, A.-L. Børresen-Dale, and C. Caldas, Breast cancer molecular stratification: from intrinsic subtypes to integrative clusters, The American Journal of Pathology 187, 2152 (2017).

[36] A. Swarbrick, A. Fernandez-Martinez, and C. M. Perou, Gene-expression profiling to decipher breast cancer inter-and intratumor heterogeneity., Cold Spring Harbor Perspectives in Medicine, a041320 (2023).

[37] D. Krijgsman, M. Hokland, and P. J. Kuppen, The role of natural killer t cells in cancer a phenotypical and functional approach, Frontiers in Immunology 9, 367 (2018).

[38] J.-S. Almeida, J. M. Casanova, M. Santos-Rosa, R. Tarazona, R. Solana, and P. Rodrigues-Santos, Natural killer t-like cells: Immunobiology and role in disease, International Journal of Molecular Sciences 24, 2743 (2023).

[39] H. Ali, E. Provenzano, S.-J. Dawson, F. Blows, B. Liu, M. Shah, H. Earl, C. Poole, L. Hiller, J. Dunn, et al., Association between cd8+ t-cell infiltration and breast cancer survival in 12 439 patients, Annals of Oncology 25, 1536 (2014).

[40] V. Sisirak, J. Faget, M. Gobert, N. Goutagny, N. Vey, I. Treilleux, S. Renaudineau, G. Poyet, S. I. Labidi-Galy, S. Goddard-Leon, et al., Impaired ifn-α production by plasmacytoid dendritic cells favors regulatory t-cell expansion that may con-tribute to breast cancer progressionpdc in breast cancer pathophysiology, Cancer Research 72, 5188 (2012).

[41] H. Lee, H. J. Lee, I. H. Song, W. S. Bang, S.-H. Heo, G. Gong, and I. A. Park, Cd11c-positive dendritic cells in triple-negative breast cancer, In Vivo 32, 1561 (2018).

[42] I. Treilleux, J.-Y. Blay, N. Bendriss-Vermare, I. Ray-Coquard, T. Bachelot, J.-P. Guastalla, A. Bremond, S. Goddard, J.-J. Pin, C. Barthelemy-Dubois, et al., Dendritic cell infiltration and prognosis of early stage breast cancer, Clinical Cancer Research 10, 7466 (2004).

[43] J. Szpor, J. Streb, A. Glajcar, P. Frączek, A. Winiarska, K. E. Tyrak, P. Basta, K. Okoń, R. Jach, and D. Hodorowicz-Zaniewska, Dendritic cells are associated with prognosis and survival in breast cancer, Diagnostics 11, 702 (2021).

[44] T. Accogli, M. Bruchard, and F. Végran, Modulation of cd4 t cell response according to tumor cytokine microenvironment, Cancers 13, 373 (2021).

[45] C. A. Huertas-Caro, M. A. Ramírez, L. Rey-Vargas, L. M. Bejarano-Rivera, D. F. Ballen, M. Nuñez, J. C. Mejía, L. F. Sua-Villegas, A. Cock-Rada, J. Zabaleta, et al., Tumor infiltrating lymphocytes (tils) are a prognosis biomarker in colombian patients with triple negative breast cancer, Scientific Reports 13, 21324 (2023).

[46] H. Matsumoto, A. A. Thike, H. Li, J. Yeong, S.-l. Koo, R. A. Dent, P. H. Tan, and J. Iqbal, Increased cd4 and cd8-positive t cell infiltrate signifies good prognosis in a subset of triple-negative breast cancer, Breast cancer research and treatment 156, 237 (2016).

[47] S. Z. Wu and A. Swarbrick, Single-cell advances in stromal-leukocyte interactions in cancer, Immunological Reviews 302, 286 (2021).

[48] J.-T. Chi, E. H. Rodriguez, Z. Wang, D. S. A. Nuyten, S. Mukherjee, M. v. de Rijn, M. J. v. de Vijver, T. Hastie, and P. O. Brown, Gene expression programs of human smooth muscle cells: tissue-specific differentiation and prognostic significance in breast cancers, PLoS genetics 3, e164 (2007).

[49] F. G. L. Mundim, F. S. Pasini, M. M. Brentani, F. A. Soares, S. Nonogaki, and A. F. L. Waitzberg, Myc is expressed in the stromal and epithelial cells of primary breast carcinoma and paired nodal metastases, Molecular and clinical oncology 3, 506 (2015).

[50] X. Tekpli, T. Lien, A. H. Røssevold, D. Nebdal, E. Borgen, H. O. Ohnstad, J. A. Kyte, J. Vallon-Christersson, M. Fon-gaard, E. U. Due, et al., An independent poor-prognosis subtype of breast cancer defined by a distinct tumor immune microenvironment, Nature Communications 10, 5499 (2019).

[51] S. Brabletz, H. Schuhwerk, T. Brabletz, and M. P. Stemmler, Dynamic emt: a multi-tool for tumor progression, The EMBO journal 40, e108647 (2021).

[52] A. Silini, C. Ghilardi, S. Figini, F. Sangalli, R. Fruscio, R. Dahse, R. B. Pedley, R. Giavazzi, and M. Bani, Regulator of g-protein signaling 5 (rgs5) protein: a novel marker of cancer vasculature elicited and sustained by the tumor’s proangiogenic microenvironment, Cellular and molecular life sciences 69, 1167 (2012).

[53] Z. Xu, Y. Zuo, J. Wang, Z. Yu, F. Peng, Y. Chen, Y. Dong, X. Hu, Q. Zhou, H. Ma, et al., Overexpression of the regulator of g-protein signaling 5 reduces the survival rate and enhances the radiation response of human lung cancer cells, Oncology reports 33, 2899 (2015).

[54] M. Oshi, M. R. Huyser, L. Le, Y. Tokumaru, L. Yan, R. Matsuyama, I. Endo, and K. Takabe, Abundance of microvascular endothelial cells is associated with response to chemotherapy and prognosis in colorectal cancer, Cancers 13, 1477 (2021).

[55] X. Wan, J. Hou, S. Liu, Y. Zhang, W. Li, Y. Zhang, and Y. Ding, Estrogen receptor α mediates doxorubicin sensitivity in breast cancer cells by regulating e-cadherin, Frontiers in Cell and Developmental Biology 9, 583572 (2021).

[56] N. P. McAndrew and R. S. Finn, Clinical review on the management of hormone receptor–positive metastatic breast cancer, JCO oncology practice 18, 319 (2022).

[57] E. Allison, S. Edirimanne, J. Matthews, and S. J. Fuller, Breast cancer survival outcomes and tumor-associated macrophage markers: a systematic review and meta-analysis, Oncology and Therapy 11, 27 (2023).

[58] S. Onkar, J. Cui, J. Zou, C. Cardello, A. R. Cillo, M. R. Uddin, A. Sagan, M. Joy, H. U. Osmanbeyoglu, K. L. Pogue-Geile, et al., Immune landscape in invasive ductal and lobular breast cancer reveals a divergent macrophage-driven microenvironment, Nature Cancer 4, 516 (2023).

[59] Y.-K. Huang, M. Wang, Y. Sun, N. Di Costanzo, C. Mitchell, A. Achuthan, J. A. Hamilton, R. A. Busuttil, and A. Boussioutas, Macrophage spatial heterogeneity in gastric cancer defined by multiplex immunohistochemistry, Nature communications 10, 3928 (2019).

[60] M. Yang, D. McKay, J. W. Pollard, and C. E. Lewis, Diverse functions of macrophages in different tumor microenvironments, Cancer research 78, 5492 (2018).

[61] I. Nasir, C. McGuinness, A. R. Poh, M. Ernst, P. K. Darcy, and K. L. Britt, Tumor macrophage functional heterogeneity can inform the development of novel cancer therapies, Trends in Immunology 44, 971 (2023).

[62] M. Matusiak, J. W. Hickey, B. Luca, G. Lu, L. Kidziński, S. Zhu, D. R. C. Colburg, D. J. Phillips, S. W. Brubaker, G. W. Charville, et al., A spatial map of human macrophage niches reveals context-dependent macrophage functions in colon and breast cancer, Research Square (2023).

[63] L. Fang, K. Liu, C. Liu, X. Wang, W. Ma, W. Xu, J. Wu, and C. Sun, Tumor accomplice: T cell exhaustion induced by chronic inflammation, Frontiers in Immunology 13, 979116 (2022).

[64] Y. Qian, Y. Yin, X. Zheng, Z. Liu, and X. Wang, Metabolic regulation of tumor-associated macrophage heterogeneity: insights into the tumor microenvironment and immunotherapeutic opportunities, Biomarker Research 12, 1 (2024).

[65] J. Wagner, M. A. Rapsomaniki, S. Chevrier, T. Anzeneder, C. Langwieder, A. Dykgers, M. Rees, A. Ramaswamy, S. Muenst, S. D. Soysal, et al., A single-cell atlas of the tumor and immune ecosystem of human breast cancer, Cell 177, 1330 (2019).

[66] B. G. Nixon, F. Kuo, L. Ji, M. Liu, K. Capistrano, M. Do, R. A. Franklin, X. Wu, E. R. Kansler, R. M. Srivastava, et al., Tumor-associated macrophages expressing the transcription factor irf8 promote t cell exhaustion in cancer, Immunity 55, 2044 (2022).

[67] T. Matsumoto, N. Takahashi, T. Kojima, Y. Yoshioka, J. Ishikawa, K. Furukawa, K. Ono, M. Sawada, N. Ishiguro, and A. Yamamoto, Soluble siglec-9 suppresses arthritis in a collagen-induced arthritis mouse model and inhibits m1 activation of raw264. 7 macrophages, Arthritis research & therapy 18, 1 (2016).

[68] P. Schmassmann, J. Roux, A. Buck, N. Tatari, S. Hogan, J. Wang, S. Lee, B. Snijder, T. A. Martins, M.-F. Ritz, et al., The siglec-sialic acid-axis is a target for innate immunotherapy of glioblastoma, bioRxiv, 2022 (2022).

[69] Y. Mei, X. Wang, J. Zhang, D. Liu, J. He, C. Huang, J. Liao, Y. Wang, Y. Feng, H. Li, et al., Siglec-9 acts as an immune-checkpoint molecule on macrophages in glioblastoma, restricting t-cell priming and immunotherapy response, Nature Cancer 4, 1273 (2023).

[70] M. Lustig, C. Chan, J. M. Jansen, M. Bräutigam, M. A. Kölling, C. L. Gehlert, N. Baumann, S. Mester, S. Foss, J. T. Andersen, et al., Disruption of the sialic acid/siglec-9 axis improves antibody-mediated neutrophil cytotoxicity towards tumor cells, Frontiers in Immunology 14, 1178817 (2023).

[71] E. L. Scolan, T. Tse, M. Krimm, W. Garner, H. Assi, J. Razo, L. Wong, K. Wong, V. Singson, J. Leong, et al., A probody drug conjugate targeting cd166 (alcam) enhances preclinical antitumor activity of a probody therapeutic targeting pd-1, Cancer Research 79, 3202 (2019).

[72] Y. Yang, A. J. Sanders, Q. P. Dou, D. G. Jiang, A. X. Li, and W. G. Jiang, The clinical and theranostic values of activated leukocyte cell adhesion molecule (alcam)/cd166 in human solid cancers, Cancers 13, 5187 (2021).

[73] I.-M. Launonen, E. P. Erkan, I. Niemiec, A. Junquera, M. Hincapie-Otero, D. Afenteva, Z. Liang, M. Salko, A. Szabo, F. Perez-Villatoro, et al., Chemotherapy induces myeloid-driven spatial t-cell exhaustion in ovarian cancer, bioRxiv (2024).

[74] C. H. Ries, M. A. Cannarile, S. Hoves, J. Benz, K. Wartha, V. Runza, F. Rey-Giraud, L. P. Pradel, F. Feuerhake Klaman, et al., Targeting tumor-associated macrophages with anti-csf-1r antibody reveals a strategy for cancer therapy, Cancer cell 25, 846 (2014).

[75] Y. N. Lamb, Pexidartinib: first approval, Drugs 79, 1805 (2019).

[76] S. Kuemmel, M. Campone, D. Loirat, R. L. Lopez, J. T. Beck, M. De Laurentiis, S.-A. Im, S.-B. Kim, A. Kwong, G. G. Steger, et al., A randomized phase ii study of anti-csf1 monoclonal antibody lacnotuzumab (mcs110) combined with gemcitabine and carboplatin in advanced triple-negative breast cancer, Clinical Cancer Research 28, 106 (2022).

[77] M. E. Fernandez, C. B. Fines, F. S. Subul, S. S. Alhudaithi, H. D. Bear, D. H. Sweet, and S. R. da Rocha, Macrophage-targeting immunotherapy for triple negative breast cancer, Cancer Research 83, 2328 (2023).

[78] Y. Komohara, D. Kurotaki, H. Tsukamoto, Y. Miyasato, H. Yano, C. Pan, Y. Yamamoto, and Y. Fujiwara, Involvement of protumor macrophages in breast cancer progression and characterization of macrophage phenotypes, Cancer Science 114, 2220 (2023).

[79] D. G. DeNardo and B. Ruffell, Macrophages as regulators of tumour immunity and immunotherapy, Nature Reviews Immunology 19, 369 (2019).

[80] Z. Duan and Y. Luo, Targeting macrophages in cancer immunotherapy, Signal transduction and targeted therapy 6, 127 (2021).

[81] S. Xu, C. Wang, L. Yang, J. Wu, M. Li, P. Xiao, Z. Xu, Y. Xu, and K. Wang, Targeting immune checkpoints on tumor-associated macrophages in tumor immunotherapy, Frontiers in Immunology 14, 1199631 (2023).

[82] P. Shahrouzi, Y. Azimzade, W. Brankiewicz, S. Bhatia, D. Kunke, D. Richard, X. Tekpli, V. N. Kristensen, and P. H. Duijf, Loss of chromosome cytoband 13q14. 2 orchestrates breast cancer pathogenesis and drug response, bioRxiv, 2024 (2024).

[83] L. Silwal-Pandit, S. Nord, H. von der Lippe Gythfeldt, E. K. Møller, T. Fleischer, E. Rødland, M. Krohn, E. Borgen, Ø. Garred, T. Olsen, et al., The longitudinal transcriptional response to neoadjuvant chemotherapy with and without bevacizumab in breast cancermolecular response in bevacizumab-treated breast tumors, Clinical Cancer Research 23, 4662 (2017).

[84] L. A. Korde, L. Lusa, L. McShane, P. F. Lebowitz, L. Lukes, K. Camphausen, J. S. Parker, S. M. Swain, K. Hunter, and J. A. Zujewski, Gene expression pathway analysis to predict response to neoadjuvant docetaxel and capecitabine for breast cancer, Breast Cancer Research and Treatment 119, 685 (2010).

[85] Y. Lin, S. Lin, M. Watson, K. M. Trinkaus, S. Kuo, M. J. Naughton, K. Weilbaecher, T. P. Fleming, and R. L. Aft, A gene expression signature that predicts the therapeutic response of the basal-like breast cancer to neoadjuvant chemotherapy, Breast Cancer Research and Treatment 123, 691 (2010).

[86] V. Popovici, W. Chen, B. D. Gallas, C. Hatzis, W. Shi, F. W. Samuelson, Y. Nikolsky, M. Tsyganova, A. Ishkin, T. Nikolskaya, et al., Effect of training-sample size and classification difficulty on the accuracy of genomic predictors, Breast Cancer Research 12, 1 (2010).

[87] A. Tabchy, V. Valero, T. Vidaurre, A. Lluch, H. Gomez, M. Martin, Y. Qi, L. J. Barajas-Figueroa, E. Souchon, C. Coutant, et al., Evaluation of a 30-gene paclitaxel, fluorouracil, doxorubicin, and cyclophosphamide chemotherapy response predictor in a multicenter randomized trial in breast cancer30-gene t/fac response predictor in breast cancer, Clinical Cancer Research 16, 5351 (2010).

[88] T. Iwamoto, G. Bianchini, D. Booser, Y. Qi, C. Coutant, C. Ya-Hui Shiang, L. Santarpia, J. Matsuoka, G. N. Hortobagyi, W. F. Symmans, et al., Gene pathways associated with prognosis and chemotherapy sensitivity in molecular subtypes of breast cancer, Journal of the National Cancer Institute 103, 264 (2011).

[89] K. Shen, N. Song, Y. Kim, C. Tian, S. D. Rice, M. J. Gabrin, W. F. Symmans, L. Pusztai, and J. K. Lee, A systematic evaluation of multi-gene predictors for the pathological response of breast cancer patients to chemotherapy, PloS One 7, e49529 (2012).

[90] S. Glück, J. S. Ross, M. Royce, E. F. McKenna, C. M. Perou, E. Avisar, and L. Wu, Tp53 genomics predict higher clinical and pathologic tumor response in operable early-stage breast cancer treated with docetaxel-capecitabine±trastuzumab, Breast Cancer Research and Treatment 132, 781 (2012).

[91] K. Shen, Y. Qi, N. Song, C. Tian, S. D. Rice, M. J. Gabrin, S. L. Brower, W. F. Symmans, J. A. O’Shaughnessy, and F. A. Holmes, Cell line derived multi-gene predictor of pathologic response to neoadjuvant chemotherapy in breast cancer: a validation study on us oncology 02-103 clinical trial, BMC Medical Genomics 5, 1 (2012).

[92] J. A. Bauer, A. B. Chakravarthy, J. M. Rosenbluth, D. Mi, E. H. Seeley, N. De Matos Granja-Ingram, M. G. Olivares, M. C. Kelley, I. A. Mayer, I. M. Meszoely, et al., Identification of markers of taxane sensitivity using proteomic and genomic analyses of breast tumors from patients receiving neoadjuvant paclitaxel and radiationmarkers of taxane sensitivity in breast cancer, Clinical Cancer Research 16, 681 (2010).

[93] C. Hatzis, L. Pusztai, V. Valero, D. J. Booser, L. Esserman, A. Lluch, T. Vidaurre, F. Holmes, E. Souchon, H. Wang, et al., A genomic predictor of response and survival following taxane-anthracycline chemotherapy for invasive breast cancer, JAMA 305, 1873 (2011).

[94] M. J. M. Magbanua, D. M. Wolf, C. Yau, S. E. Davis, J. Crothers, A. Au, C. M. Haqq, C. Livasy, H. S. Rugo, L. Esserman, et al., Serial expression analysis of breast tumors during neoadjuvant chemotherapy reveals changes in cell cycle and immune pathways associated with recurrence and response, Breast Cancer Research 17, 1 (2015).

[95] T. Miyake, T. Nakayama, Y. Naoi, N. Yamamoto, Y. Otani, S. J. Kim, K. Shimazu, A. Shimomura, N. Maruyama, Y. Tamaki, et al., Gstp 1 expression predicts poor pathological complete response to neoadjuvant chemotherapy in er-negative breast cancer, Cancer Science 103, 913 (2012).

[96] J. C. Liu, V. Voisin, G. D. Bader, T. Deng, L. Pusztai, W. F. Symmans, F. J. Esteva, S. E. Egan, and E. Zacksenhaus, Seventeen-gene signature from enriched her2/neu mammary tumor-initiating cells predicts clinical outcome for human her2+: Erα-breast cancer, Proceedings of the National Academy of Sciences 109, 5832 (2012).

[97] A. Prat, G. Bianchini, M. Thomas, A. Belousov, M. C. Cheang, A. Koehler, P. Gómez, V. Semiglazov, W. Eiermann, S. Tjulandin, et al., based pam50 subtype predictor identifies higher responses and improved survival outcomes in her2-positive breast cancer in the noah study, Clinical Cancer Research 20, 511 (2014).

[98] F. Pedregosa, G. Varoquaux, A. Gramfort, V. Michel, B. Thirion, O. Grisel, M. Blondel, P. Prettenhofer, R. Weiss, V. Dubourg, J. Vanderplas, A. Passos, D. Cournapeau, M. Brucher, M. Perrot, and E. Duchesnay, Scikit-learn: Machine learning in Python, Journal of Machine Learning Research 12, 2825 (2011).

[99] D. Meyer and C. Buchta, proxy: Distance and similarity measures. r package version 0. 4–15 (2015).

[100] T. M. Therneau, A Package for Survival Analysis in R, (2024).

[101] C. Zhang, Q. Sheng, X. Zhang, K. Xu, X. Jin, W. Zhou, M. Zhang, D. Lv, C. Yang, Y. Li, et al., Prioritizing exhausted t cell marker genes highlights immune subtypes in pan-cancer, Iscience 26, 106484 (2023).

