## Supplementary Info for "Integrated multiomics analysis unveils how macrophages drive immune suppression in breast tumors and affect clinical outcomes"

(Dated: November 9, 2024)

### I. DECONVOLUTION

Deconvolution of bulk gene expression or RNA-seq data is the process of estimating the proportions of different cell types that contribute to the overall measurement. Assuming the final measurement is a linear combination of the expressions from various cell types, it can be written as [1]:

$$\mathbf{P}_j = \mathbf{SM} \cdot \boldsymbol{\beta}_j + \boldsymbol{\epsilon} \tag{1}$$

In this equation, $\mathbf{P}j$ represents the measured expression/RNA-seq values from tumor sample $j$, $\mathbf{SM}$ is the signature matrix, $\boldsymbol{\beta}j$ is a vector of mixing fractions for sample $j$, and $\boldsymbol{\epsilon}$ represents the noise. For $Pij$, the expression of gene $i$ in sample $j$, we have: $Pij = \sum_k SM_{ik}\beta_{kj}$, where $SM_{ik}$ is the average expression of gene $i$ in cell type $k$ and $\beta_{kj}$ is the fraction of cell type $k$ in sample $j$.

We used 10 signature matrices from our previous work [2], created using high-resolution scRNA-seq data from [3]. To estimate cell fractions, we used a common method called CIBERSORTx [4]. Bulk RNA-seq data from TCGA and GEPs data from MBRC were downloaded from cBioPortal. TCGA data were log2-normalized and scaled, while MBRC data were scaled. We estimated cell fractions using either the "Impute Cell Fractions" module of the CIBERSORTx webtool with default settings or the Docker version of CIBERSORTx with similar settings. We then averaged the results from all 10 estimates.

### II. TME COMPOSITION ACROSS ER SUBTYPES

We compared the fraction of cell types across ER+ and ER- samples in both TCGA and MBRC, revealing consistent differences in the fraction of cancer and immune cells, as shown in Fig. S1.

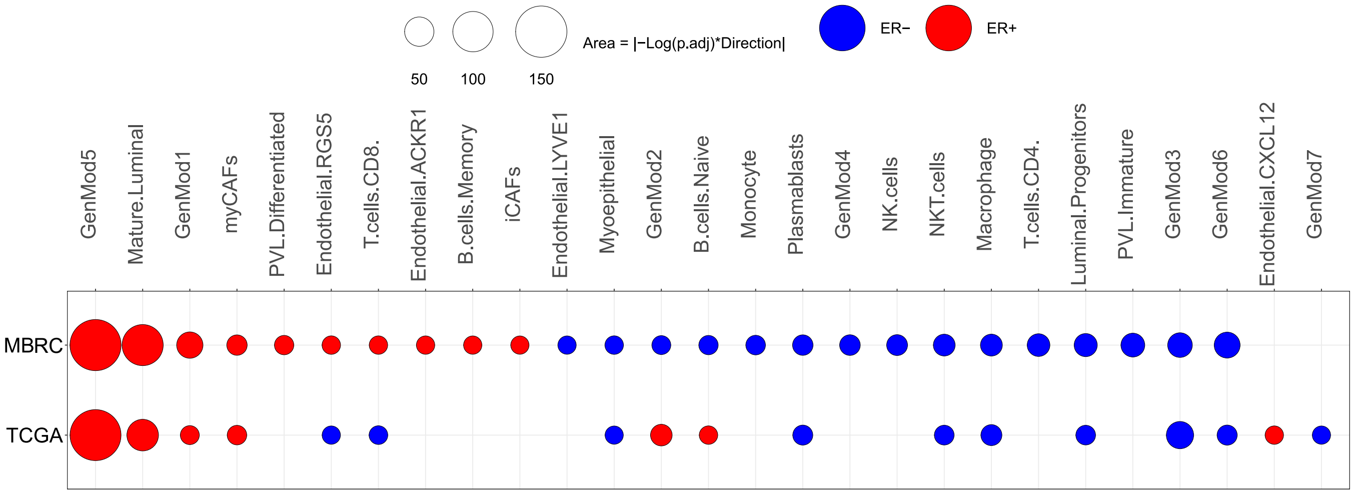


(a)

Figure S1: Comparison of cell type fractions in ER+ vs. ER- samples.

### III. SURVIVAL SCORES IN BREAST CANCER SUBTYPES

ER and PAM50 subtypes were analyzed using the same pipeline to calculate Survival Scores. Fig. S2 shows the results for all subtypes in TCGA and MBRC separately.

### A. Survival Scores in ER subtypes

**Cell types with positive Survival Scores in ER+:** Among immune cells, NKT cells, CD8 T cells, and DCs have positive Survival Scores. In cancer cells, GenMod5 and GenMod2 show positive Survival Scores. From normal epithelial cells, ACKR1 endothelials and cancer-associated fibroblasts with inflammatory features (iCAFs) show positive Survival Scores.

**Cell types with negative Survival Scores in ER+:** RGS5 endothelials and differentiated PVLs from microvascular cells have negative Survival Scores. Among immune cells, NK cells and macrophages show negative Survival Scores. GenMod3 from cancer cells, myCAFs from stromal cells, and luminal progenitors from normal epithelial cells also show negative Survival Scores.

**Cell types with positive Survival Scores in ER-:** NKT cells, plasmablasts, CD4 T cells, and naive B cells, all immune cells, show positive Survival Scores.

**Cell types with negative Survival Scores in ER-:** GenMod6 and luminal progenitors show negative Survival Scores.

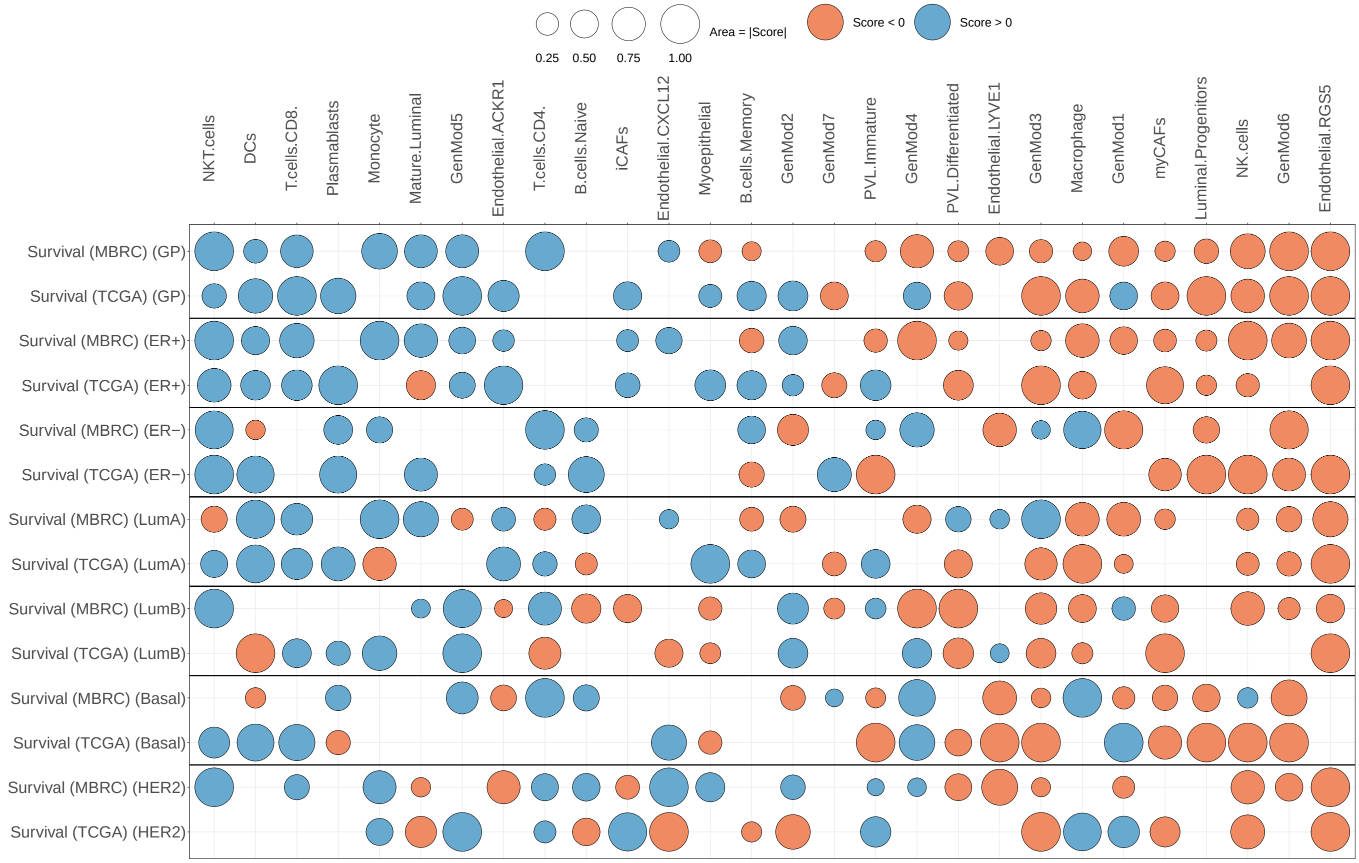


Figure S2: Survival Scores for subtypes in MBRC and TCGA datasets.

### B. Survival Scores in PAM50 subtypes

CD8 T cells, DCs, and ACKR1 endothelials show positive Survival Scores in LumA tumors. GenMod1, macrophages, GenMod6, NK cells, and RGS5 endothelials show negative Survival Scores in LumA. GenMod5 and GenMod2 show positive Survival Scores in LumB tumors. Myoepithelials, macrophages, differentiated PVLs, GenMod3, myCAFs, and RGS5 endothelials show negative Survival Scores in LumB. GenMod4 shows positive Survival Scores in Basal. Luminal progenitors, LYVE1 endothelials, GenMod3, GenMod6, and myCAFs show negative Survival Scores in Basal. Monocytes, CD4 T cells, and immature PVLs show positive Survival Scores in HER2. Mature luminals, NK cells, and RGS5 endothelials show negative Survival Scores in HER2.

### IV. ALIGNMENT OF PCR AND SURVIVAL

Generally, in the breast cancer population, ER+ tumors have a lower pCR rate but better survival compared to ER- tumors. This well-established fact in breast tumors highlights the importance of evaluating the relationship between pCR and survival. To quantify this alignment, we developed an intuitive method to explore the alignment between pCR and survival from a TME perspective. We defined and calculated an "Alignment Score" as $\sum_{i=1}^{28} \text{pCR}_i \cdot \text{Survival}_i$, where $\text{pCR}_i$ and $\text{Survival}_i$ are the corresponding scores for cell type $i$.

We calculated the Alignment Score separately for TCGA and MBRC datasets, as shown in Fig. S3. In the GP, as mentioned earlier, pCR does not imply better survival. This is evident in the Alignment Score of GP for both datasets. Within ER subtypes, ER- shows positive alignment, suggesting a positive association between these two clinical outcomes. For ER+, the results are inconsistent. Interestingly, PAM50 subtypes show a clear positive or negative alignment. Both LumA and LumB show negative Alignment Scores across both TCGA and MBRC. HER2 and Basal subtypes show positive Alignment Scores, following the ER- results.

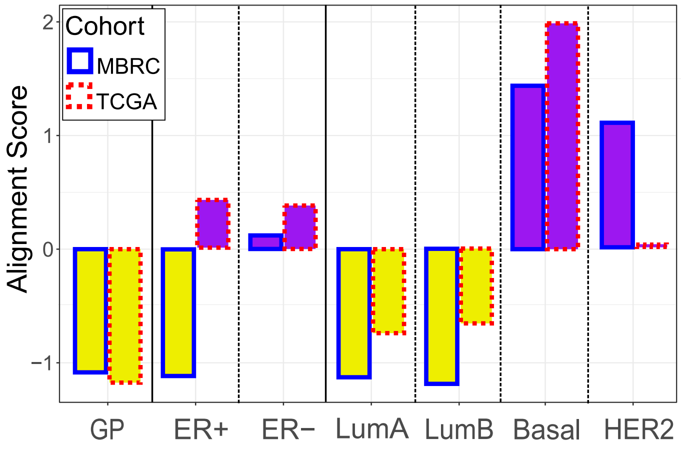


Figure S3: Alignment Score calculated for all groups.

### V. GENMOD5 FRACTION STRATIFIES BREAST TUMORS INTO GROUPS WITH DISTINCT CLINICAL OUTCOMES

A possible explanation for the negative Alignment Score in LumB tumors is that there might be different subtypes within this group with unique pCR rates and survival outcomes. Looking back at Fig. **??** in the main text, we explored if the fraction of GenMod5 can stratify LumB samples. Therefore, we divided LumB samples into three groups with equal numbers of samples based on the density of GenMod5 (Low, Int, and High). We first selected the NAC cohort, which includes 464 samples. Divided into three groups, we separated the group with a higher fraction of GenMod5 and compared their pCR rate with the other two groups, which were combined into a single group "Low/Int". As expected, the pCR rate is less than 45% of that of the "Low/Int" group, as shown in Fig. S4(a). Interestingly, the risk of recurrence (RoR) score [5] for the same group is significantly lower (refer to Fig. S4(b)), suggesting the existence of a subset of LumB tumors with a low pCR rate but better survival. To further explore this observation, we divided LumB samples in the MBRC and TCGA datasets into two groups based on the fraction of GenMod5 and compared their survival using a Kaplan-Meier graph. In both datasets, their survival is significantly different based on the p-value of the log-rank test, as shown in Fig. S4(c) and (d). Similarly, the GenMod5 fraction can be used to divide ER+ or GP samples into two groups with distinct behaviors, as shown in the rest of Fig. S4.

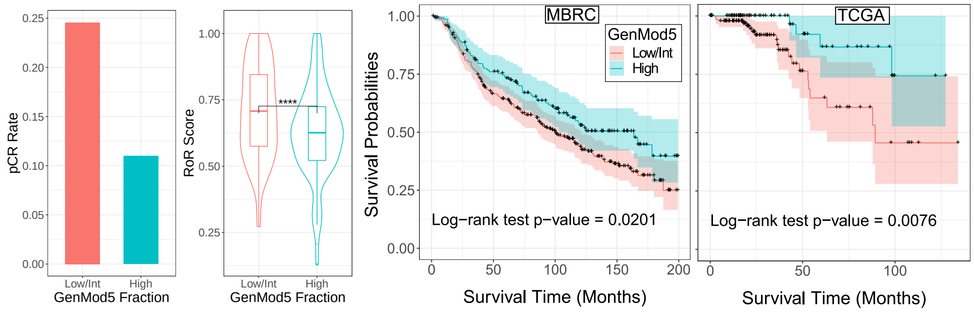


(a) NAC (LumB) (b) NAC (LumB) (c) MBRC (LumB) (d) TCGA (LumB)

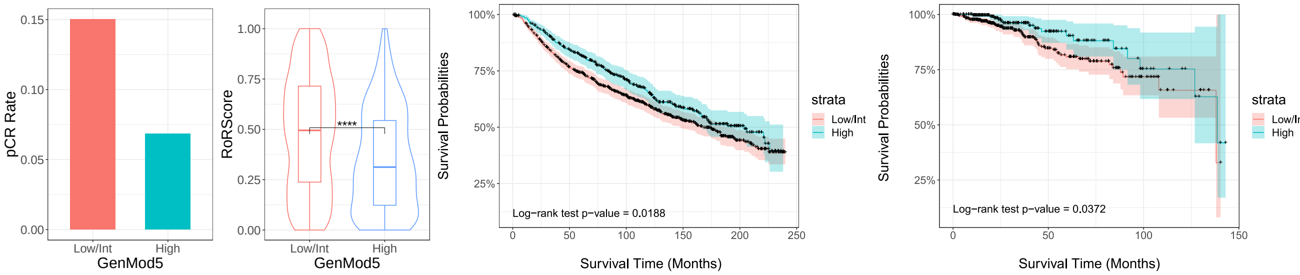


(e) NAC (ER+) (f) NAC (ER+) (g) MBRC (ER+) (h) TCGA (ER+)

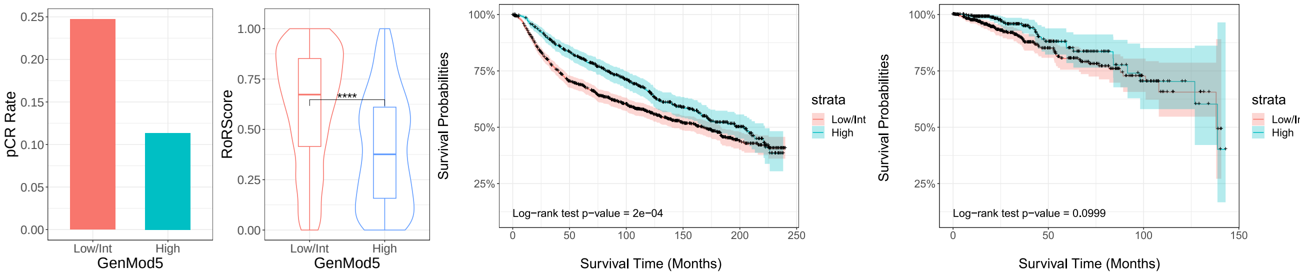


(i) NAC (GP) (j) NAC (GP) (k) MBRC (GP) (l) TCGA (GP)

Figure S4: GenMod5 fraction stratifies breast tumors into groups with distinct clinical outcomes in LumB, ER+, and GP tumors.

### VI. MACROPHAGES FRACTION STRATIFIES BREAST CANCER SAMPLES INTO GROUPS WITH DISTINCT CLINICAL OUTCOMES

Following the observation in the main text, we also divided ER+ and GP samples into two groups based on the fraction of macrophages and explored their characteristics.

**Deconvolution:** For the NAC cohort, we compared pCR rates and RoR scores for these groups of samples across ER+ and GP tumors. Following the observation for LumA samples, samples with high macrophage fraction show higher pCR rates while also having significantly higher RoR scores, as shown in Fig. S5 (a), (b), (f), and (g). In MBRC, samples with a high fraction of macrophages show significantly lower survival probabilities, as shown in Fig. S5 (c) and (h). In TCGA, however, such significance was not observed. The lack of such observation does not undermine the role of macrophages in the survival probability of the TCGA dataset, as we consistently recovered negative Survival Scores in LumA, LumB, ER+, and GP in this dataset.

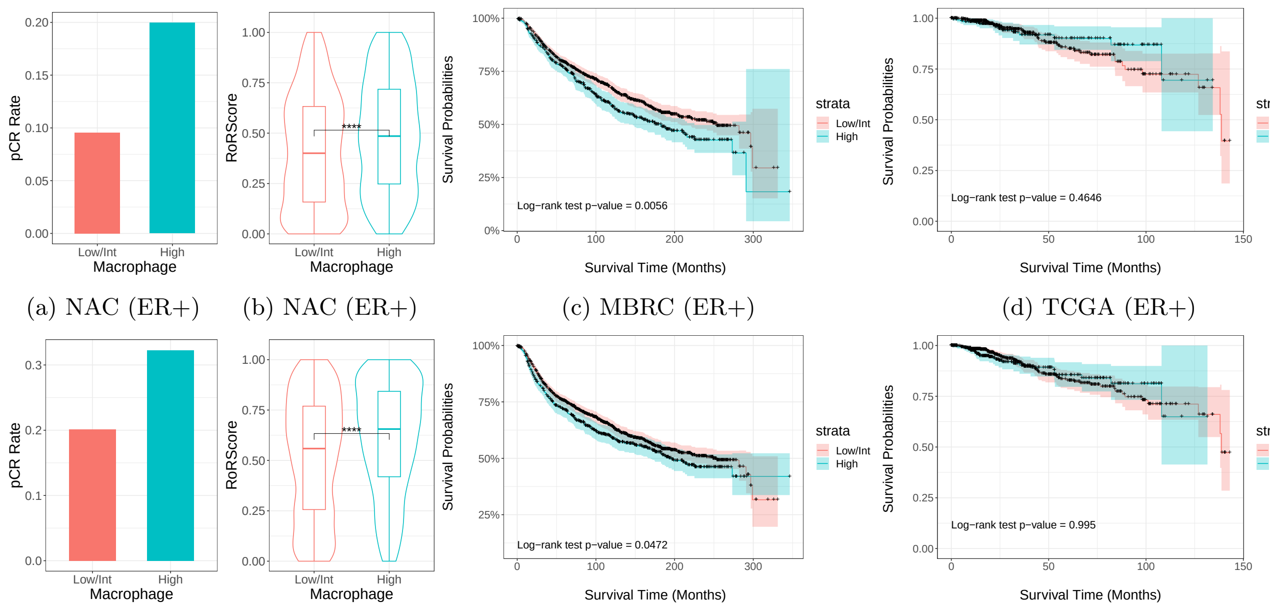


(a) NAC (ER+) (b) NAC (ER+) (c) MBRC (ER+) (d) TCGA (ER+)

(e) NAC (GP) (f) NAC (GP) (g) MBRC (GP) (h) TCGA (GP)

Figure S5: Estimated macrophage fraction from deconvolution stratifies breast cancer samples into groups with distinct clinical outcomes.

**IMC Data:** Similar to deconvolution data, we used the fraction of macrophages obtained from IMC data [6] to divide ER+ and GP samples into two groups following the same method. As Figs. S6 (a) and (b) show, in both cases, samples with a high fraction of macrophages show significantly lower survival probabilities.

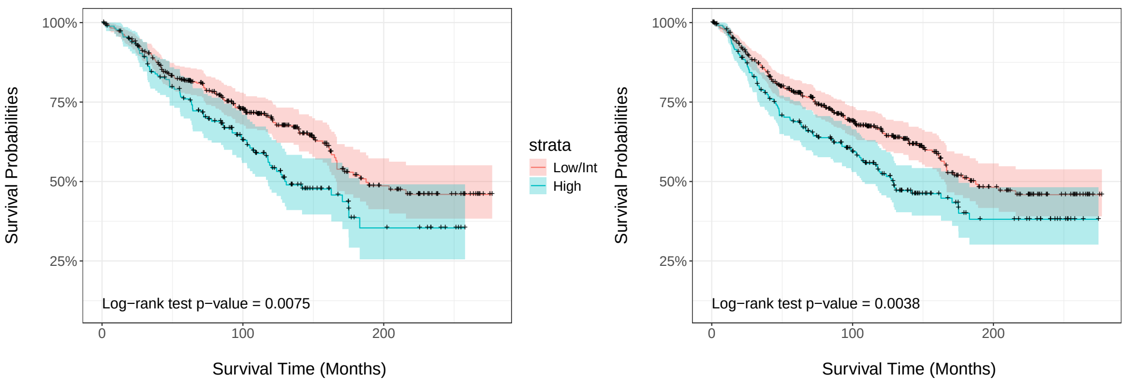


(a) ER+ (b) GP

Figure S6: Macrophages fraction in IMC data stratifies breast cancer samples in IMC data into groups with distinct clinical outcomes.

### VII. DISTANCE FROM MHC I & II$^{\mathrm{hi}}$ ECS REVERSES ROLE OF MACROPHAGES IN RFS

In LumB and ER+ samples, we divided macrophages into two groups based on their minimum distance to MHC I & II$^{\mathrm{hi}}$ ECs. The threshold distance, $r_T$, is the median of the minimum distances, $r_m$, between macrophages and the target cell type across LumB and ER+ samples, respectively. We then fit a Cox proportional hazards model [7] to each subset of macrophages. As the survival area plot [8] in Fig. S7 shows, for both LumB and ER+ samples, a higher frequency of macrophages close to (distant from) MHC I & II$^{\mathrm{hi}}$ leads to lower (higher) survival probability.

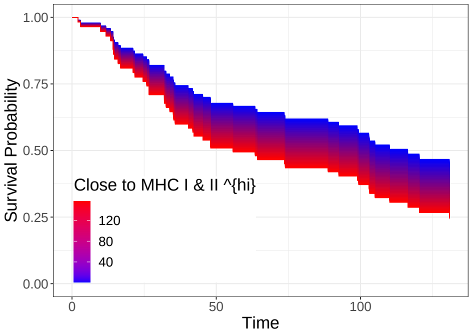


(a) LumB

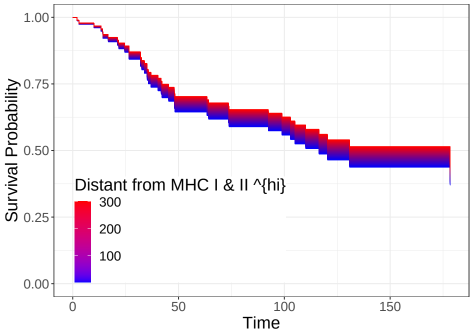


(b) LumB

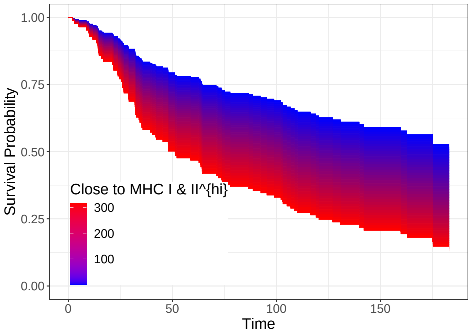


(c) ER+

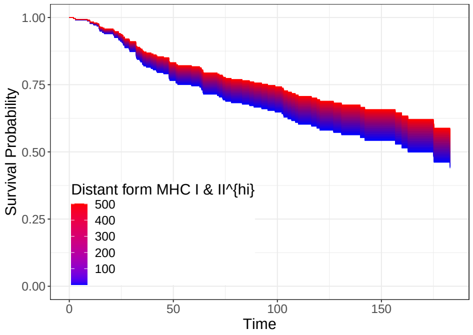


(d) ER+

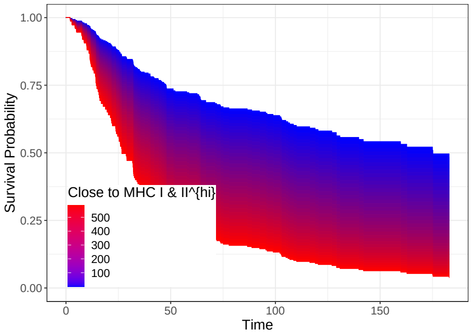


(e) GP

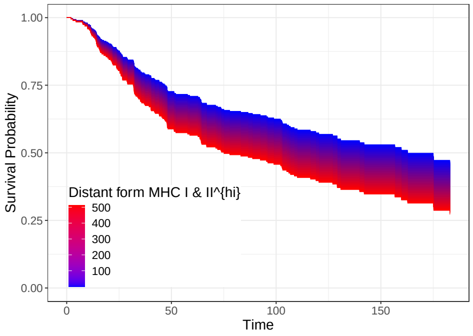


(f) GP

Figure S7: Survival area plot for the effect of the frequency of macrophages close to MHC I & II$^{\text{hi}}$ and distant from MHC I & II$^{\text{hi}}$ in LumB, ER+, and GP samples.

### VIII. M2 MACROPHAGES DIVIDED BASED ON HLA-ABC LEVEL

In treatment-naive samples from [9], which belong to triple-negative breast cancer (TNBC), we divided (M2) macrophages (the only cell type distinctly annotated as macrophages) into two groups based on their HLA-ABC levels and explored the correlation of each group with TReg and TEx. As shown in Fig. S8 (a), only HLA-ABC$^{\text{hi}}$ macrophages exhibit a significant correlation with TEx. While the correlation of HLA-ABC$^{\text{lo}}$ macrophages with TReg is significant, it is lower than that of HLA-ABC$^{\text{hi}}$ with $T_{\text{Reg}}$.

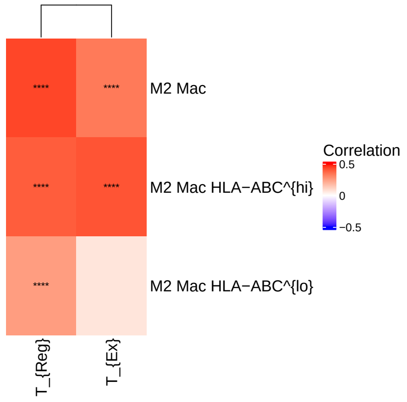


Figure S8: Correlation of macrophages subsets with TReg and TEx.

### IX. HLA-ABC$^{hi}$ ECS AND IMMUNOSUPPRESSIVE TME

To investigate whether the frequency of HLA-ABC$^{hi}$ ECs is associated with the frequency of TEx and TReg cells [10], we divided the ECs in scRNA-seq data into two groups, HLA-ABC$^{lo}$ and HLA-ABC$^{hi}$ ECs, and calculated the correlation of both groups with these T cells. However, as shown in Fig. S9, there is no significant correlation between the frequencies of EC subsets and the frequencies of TEx and TReg cells.

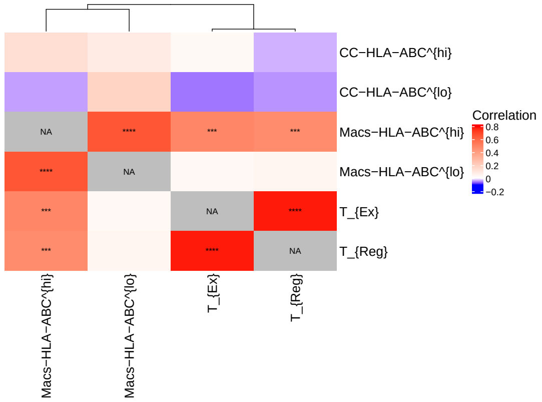


Figure S9: Correlation of HLA-ABC$^{hi/lo}$ ECs and macrophages with TReg and TEx.

### X. CELL-CELL COMMUNICATION ASSAY

Computational cell-cell communication assays are used to study and predict interactions between different cell types within a tissue or organism. These assays help understand how cells influence each other's behavior through signaling molecules. NicheNet [11] is a method that predicts ligand–target links between interacting cells by combining their expression data with prior knowledge of signaling and gene regulatory networks.

For interaction between TEx and macrophages, TEx were set as sender cells while macrophages were set as receiver cells. To run NicheNet we again followed ligand activity geneset vignette. We selected the top 50 most variant genes on macrophages as target genes of interest in receiver cells.

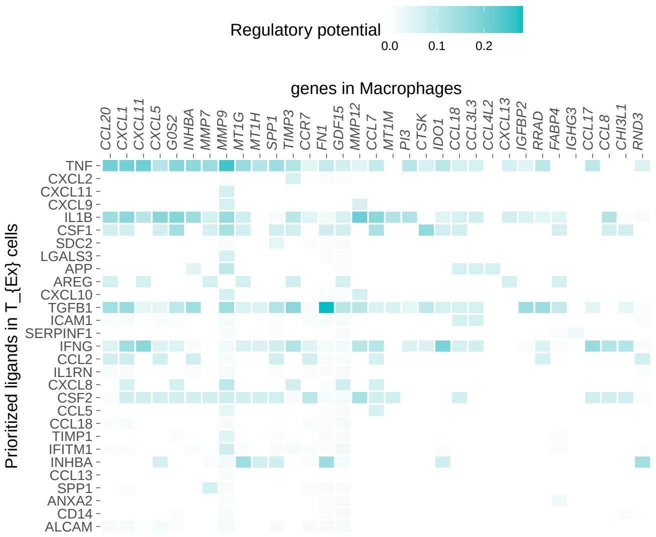


Figure S10: Interaction of $T_{Ex}$ and macrophages.

### XI. HLA-ABC AND OTHER GENES IN MACROPHAGES

We also explored the correlation between the level of HLA-ABC and some other markers that were previously shown to be involved in immunosuppressive macrophages. In particular, we plotted HLA-ABC levels vs. PDL1 and IRF8. As Figs. S11 (a) and (b) show, both of these markers are highly correlated with HLA-ABC.

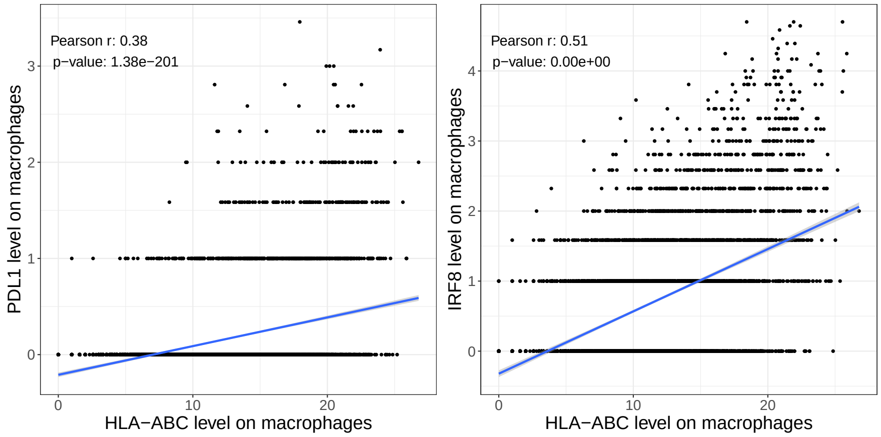


(a) (b)

Figure S11: a) Correlation of HLA-ABC vs PDL1 on macrophages in scRNA-seq. b) IRF8 vs HLA-ABC on macrophages in scRNA-seq data.


[1] F. Avila Cobos, J. Vandesompele, P. Mestdagh, and K. De Preter, Computational deconvolution of transcriptomics data from mixed cell populations, Bioinformatics **34**, 1969 (2018).

[2] Y. Azimzade, M. Haugen, X. Tekpli, C. B. Steen, T. Fleischer, D. Kilburn, H. Ma, E. V. Egeland, G. Mills, O. Engebraaten, *et al.*, Explainable machine learning reveals the role of the breast tumor microenvironment in neoadjuvant chemotherapy outcome, bioRxiv , 2023 (2023).

[3] S. Z. Wu, G. Al-Eryani, D. L. Roden, S. Junankar, K. Harvey, A. Andersson, A. Thennavan, C. Wang, J. R. Torpy, N. Bartonicek, *et al.*, A single-cell and spatially resolved atlas of human breast cancers, Nature Genetics **53**, 1334 (2021).

[4] A. M. Newman, C. B. Steen, C. L. Liu, A. J. Gentles, A. A. Chaudhuri, F. Scherer, M. S. Khodadoust, M. S. Esfahani, B. A. Luca, D. Steiner, *et al.*, Determining cell type abundance and expression from bulk tissues with digital cytometry, Nature Biotechnology **37**, 773 (2019).

[5] J. S. Parker, M. Mullins, M. C. Cheang, S. Leung, D. Voduc, T. Vickery, S. Davies, C. Fauron, X. He, Z. Hu, *et al.*, Supervised risk predictor of breast cancer based on intrinsic subtypes, Journal of Clinical Oncology **27**, 1160 (2009).

[6] E. Danenberg, H. Bardwell, V. R. Zanotelli, E. Provenzano, S.-F. Chin, O. M. Rueda, A. Green, E. Rakha, S. Aparicio, I. O. Ellis, *et al.*, Breast tumor microenvironment structures are associated with genomic features and clinical outcome, Nature Genetics **54**, 660 (2022).

[7] J. Fox and S. Weisberg, Cox proportional-hazards regression for survival data, An R and S-PLUS companion to applied regression **2002** (2002).

[8] R. Denz and N. Timmesfeld, Visualizing the (causal) effect of a continuous variable on a time-to-event outcome, Epidemiology **34**, 652 (2023).

[9] X. Q. Wang, E. Danenberg, C.-S. Huang, D. Egle, M. Callari, B. Bermejo, M. Dugo, C. Zamagni, M. Thill, A. Anton, *et al.*, Spatial predictors of immunotherapy response in triple-negative breast cancer, Nature **621**, 868 (2023).

[10] S. Tietscher, J. Wagner, T. Anzeneder, C. Langwieder, M. Rees, B. Sobottka, N. de Souza, and B. Bodenmiller, A comprehensive single-cell map of t cell exhaustion-associated immune environments in human breast cancer, Nature Communications **14**, 98 (2023).

[11] R. Browaeys, W. Saelens, and Y. Saeys, Nichenet: modeling intercellular communication by linking ligands to target genes, Nature Methods **17**, 159 (2020).
